# The fecal microbiota of lactating Holstein dairy cows: a meta-analysis highlighting key microbial profiles and methodological challenges

**DOI:** 10.1101/2025.07.21.666008

**Authors:** Lisa Arnalot, Géraldine Pascal, Sébastien Dejean, Gilles Foucras, Asma Zened

## Abstract

In cattle, the digestive microbiota has been poorly studied outside the rumen, despite its recognized role in animal physiology, health, and production. This study aimed to provide a comprehensive meta-analysis of the fecal microbiota of lactating Holstein dairy cows. A systematic online search of PubMed was performed in June 2023 to identify studies on the fecal microbiota of lactating Holstein dairy cows. Of the 526 articles retrieved from the systematic online search, only 28 met our required inclusion criteria. Raw sequencing data targeting the 16S rRNA gene (V3-V4 region) were obtained either from public repositories or directly from the authors. Two recently published articles were included as they met the inclusion criteria. A total of 2,136 samples were included in the analysis. The number of sequences per sample varied considerably between studies, and the metadata were sparse. The core microbiota was identified as the most prevalent and shared taxa, comprising 21 microbiota families that accounted for 82% of the relative abundance.

The observed clustering, which depended on study, highlighted the significant impact of environmental factors and experimental conditions on the microbial communities of cattle. A multi-group analysis was performed to correct for the study effect and successfully identified three microbiota profiles.

The meta-analysis approach is a rigorous and systematic method of analyzing research findings that allows for the generation of reliable and reproducible results. This approach ensures the independence of results from a single experimental facility or condition, thereby enhancing the reliability and generalizability of the findings.

**Importance:** This meta-analysis provides the most comprehensive and up-to-date overview of the fecal microbiota of lactating Holstein dairy cows, based on a rigorous selection of over 2,000 samples. By controlling for inter-study variation, it successfully identified consistent microbial patterns and establishes a robust core microbiota. This work not only highlighted the microbial taxa most likely essential to bovine physiology and health, but also emphasized the need for standardized methodologies and improved data-sharing practices across studies. Ultimately, the findings pave the way for strategies that target the modulation of microbiota with potential benefits for animal health and environmental impacts.

## INTRODUCTION

Over the last 10 years, numerous studies have linked the gastrointestinal microbiota to health and specific disease conditions across species (Fan & Pedersen, 2021; O’Hara et al., 2020). In humans, mice and pigs, the digestive microbiota is typically characterized through fecal sampling (Mishra et al., 2023; Seyoum et al., 2024; Xiao et al., 2016). Conversely, most research conducted on the microbiota of the bovine digestive tract has focused on the forestomach microbiota. This topic has been extensively reviewed and is the subject of numerous recent meta-analyses, with linked to various topics, such as methane emissions, nutrition and the functions of the rumen epithelium (Ma et al., 2025; Plaizier et al., 2021; Susanto et al., 2024).

Surprisingly, the fecal microbiota of dairy cows has not been studied as thoroughly. We could only identify one meta-analysis that focused specifically on feces (Kim & Wells, 2016). More recently, Holman & Gzyl (2019) compared ruminal and fecal microbiota, including data from diverse breeds, types of cattle at different ages and physiological stages. The authors concluded that substantial variability was caused by the hypervariable region studied, the sampling location within the gastrointestinal tract and the study of origin.

The fecal microbiota of dairy cows is perhaps of greater interest than the ruminal microbiota, because samples are easier to collect, with fewer ethical and technical constraints. This makes it a more reliable source of information that could improve cow’s production traits or health, resulting in more robust and important findings (Monteiro et al., 2022, 2024; Plaizier et al., 2017; G. Zhang et al., 2019). This reinforces the importance of studying its composition and central core, which was first characterized over a decade ago as a recurrent and shared microbial community among individuals (Petri et al., 2013).

Various cattle breeds are reared on all continents, but one breed has gained importance in almost all countries with dairying in the last fifty years. The Holstein breed has indeed established itself as the most productive dairy cow worldwide. With more than 25 million heads of cattle, this breed is primarily raised for milk production, in zero-grazing or pasture conditions. Thanks to intensive genetic selection, the Holstein breed is arguably the most suitable for milk production. The present study aimed to provide an overview of the fecal microbiota of the Holstein dairy breed, which is the most important category of cattle worldwide. We examined data collected during the lactation physiological stage. Raw data from studies that sequenced the V3-V4 regions of the 16S rRNA gene were gathered to minimize variability and draw robust conclusions by using the most published hypervariable region in bovine fecal microbiota. By alleviating biases related to the study, we described the most common and abundant taxa, approaching a definition of the core microbiota of dairy cattle.

## MATERIALS AND METHODS

This work follows the Preferred Reporting Items for Systematic reviews and Meta-analyses (PRISMA) statement (Page et al., 2021).

### Literature search

Studies were identified through a systematic online search on PubMed (https://pubmed.ncbi.nlm.nih.gov/) using the following search query: “(Microbiota OR Microbiome) AND (metagenomics OR 16S sequencing) AND (intestinal OR fecal) AND (Ruminant OR dairy cattle OR cow OR bovine)”. The search was conducted on June 15, 2023. Subsequently, from June to December 2023, a Google Scholar alert (https://scholar.google.com/) was implemented to identify new publications meeting these criteria.

### Study selection

To be included in the meta-analysis, the studies had to fit the following inclusion criteria:

I. investigated the fecal microbiota of female cows.
II. analyzed the V3-V4 region of the 16S rRNA gene, as this region has been the subject of the most extensive research.
III. investigated lactating Holstein dairy cows. However, studies including other breeds were also accepted as long as the Holstein breed could be identified.
IV. include original raw data, excluding reviews or any kind of articles using data from other articles
V. The raw data of the microbiota analysis were available in a public repository: ENA (European Nucleotide Archive: https://www.ebi.ac.uk/ena), NCBI (National Center for Biotechnology Information: https://www.ncbi.nlm.nih.gov/), DDBJ (DNA Data Bank of Japan: https://www.ddbj.nig.ac.jp/). Alternatively, they could be obtained from the authors who were contacted.

Research articles and notes that met with the above criteria, were included in the analysis. Two larger datasets published in 2024 (Brulin et al., 2024) and 2025 (Arnalot et al., *Animal Microbiome*, accepted for publication) were also included in the analysis.

### Data extraction

The raw sequences were obtained directly from the aforementioned databases. The authors of articles that met the inclusion criteria were contacted to request the raw data, further information on the materials and methods used in their study, or more details related to the cows. The following information was collected, where available or kindly provided by the authors:

- Research topic: feed, health or microbiota description (including aging, transition period and microbiota stability).
- Type of farm: commercial, experimental or demonstration.
- DNA extraction method and commercial kit information, if applicable.
- Sequencing platform
- For individual cows, when applicable, whether the animal was a control or treated/diseased animal, if applicable, and other zootechnical information such as parity and/or lactation stage. Any missing or unclear information was noted as “NA” for Not Available.

### Bioinformatic processing

The raw sequences were analyzed using the FROGS v4.1.0 software (Escudié et al., 2018). We successively used *preprocessing*, *clustering*, *remove_chimera*, *cluster_filter*, *taxonomic_affiliation* tools with the parameters shown in **Supplementary Table 1** for samples processed only with read 1, due to inability to contig reads some or for all of the samples from the articles (**Table 1**), and in **Supplementary Table 2** for the others.

**Table 1:**
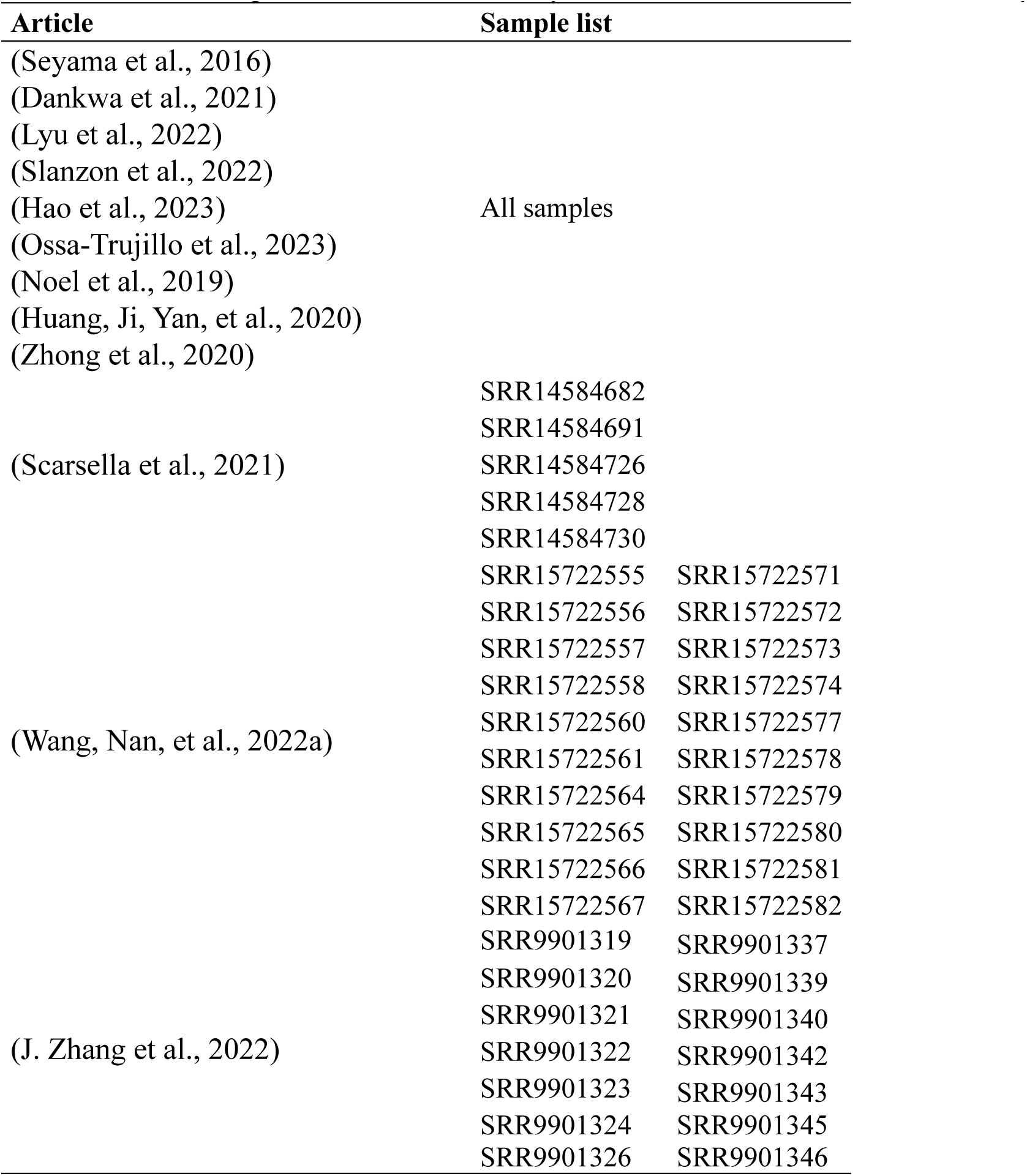
List of samples extracted where only read1 was used for downstream analysis

Three samples from Huang et al., (2020) were processed separately because they could not be properly merged with the other datasets (SRR11617225, SRR11617226 and SRR11617227). Five samples were neither processed nor used in the analysis of the data of Scarsella et al., (2021): SRR14584731, SRR14584692 and SRR14584693 had an unequal number of read1 and read2, while SRR14584729 and SRR14584631 were excluded due to atypical read lengths that differed from the other samples. Two samples (SRR9901317 and SRR9901341) from Zhang et al., (2022), three samples (SRR26046274, SRR26046275 and SRR26046276) from Li et al., (2023), and six samples (SRR12183056, SRR12183065, SRR12183074, SRR12183077, SRR12183096 and SRR12183098) from Dankwa et al., (2021) were also excluded from the analysis, as no sequences were obtained after processing and filtering.

Each bioinformatic process generated a phyloseq object (McMurdie & Holmes, 2013).

### Statistical analysis

Statistical analysis was performed using the R software, version 4.3.3 (R Core Team, 2025). First, the generated phyloseq objects were merged based on taxonomic affiliation at the genus level. When the samples were merged, NA counts were replaced with 0.

As our focus was on bacteria, clusters allocated to kingdoms other than bacteria and unclassified bacterial phyla were removed. Following the approach of Holman and Gzyl. (2019), samples containing fewer than 1,000 sequences were excluded from further analysis. The diversity indicators (observed richness and Bray-Curtis) were described with the *vegan* package (Oksanen et al., 2022). Noise removal was performed by filtering out clusters that contributed less than 1% to the total sum. The *microbiome* package was used in order to define the core microbiota (Lahti & Shetty, 2018), which was present in a minimum of 90% of the samples, as previously described by Holman et al. (2017).

The *mixOmics* package was used to perform a Principal Component Analysis (PCA) after Centered Log-Ratio (CLR) transformation and for multi-group integration (function *mint.pca*) which is designed to identify reproducible molecular signatures across different datasets, in order to correct for the article effect (Rohart, Eslami, et al., 2017; Rohart, Gautier, et al., 2017). Partition around medoid clustering was performed using the *cluster* package (Maechler, 2018), to determine the microbiota profile. The *ancombc2* from the package ANCOM-BC (prvcut =0.5) was used in order to identify differential taxa across the microbiota profiles (Lin & Peddada, 2020).

## RESULTS

### Selection criteria and the results of the literature search

A total of 28 articles were selected as fitting the required inclusion criteria (see **Figure 1** for the selection process), along with two additional recently published datasets from our institute. The articles are presented in **Supplementary Table 3**. All the articles contained samples from individual cows, except for those by Ossa-Trujillo et al., (2023) which included pooled fecal samples from lactating Holstein cows, and by Sun et al., (2017) which represented mixed samples from the same cow.

**Figure 1:**
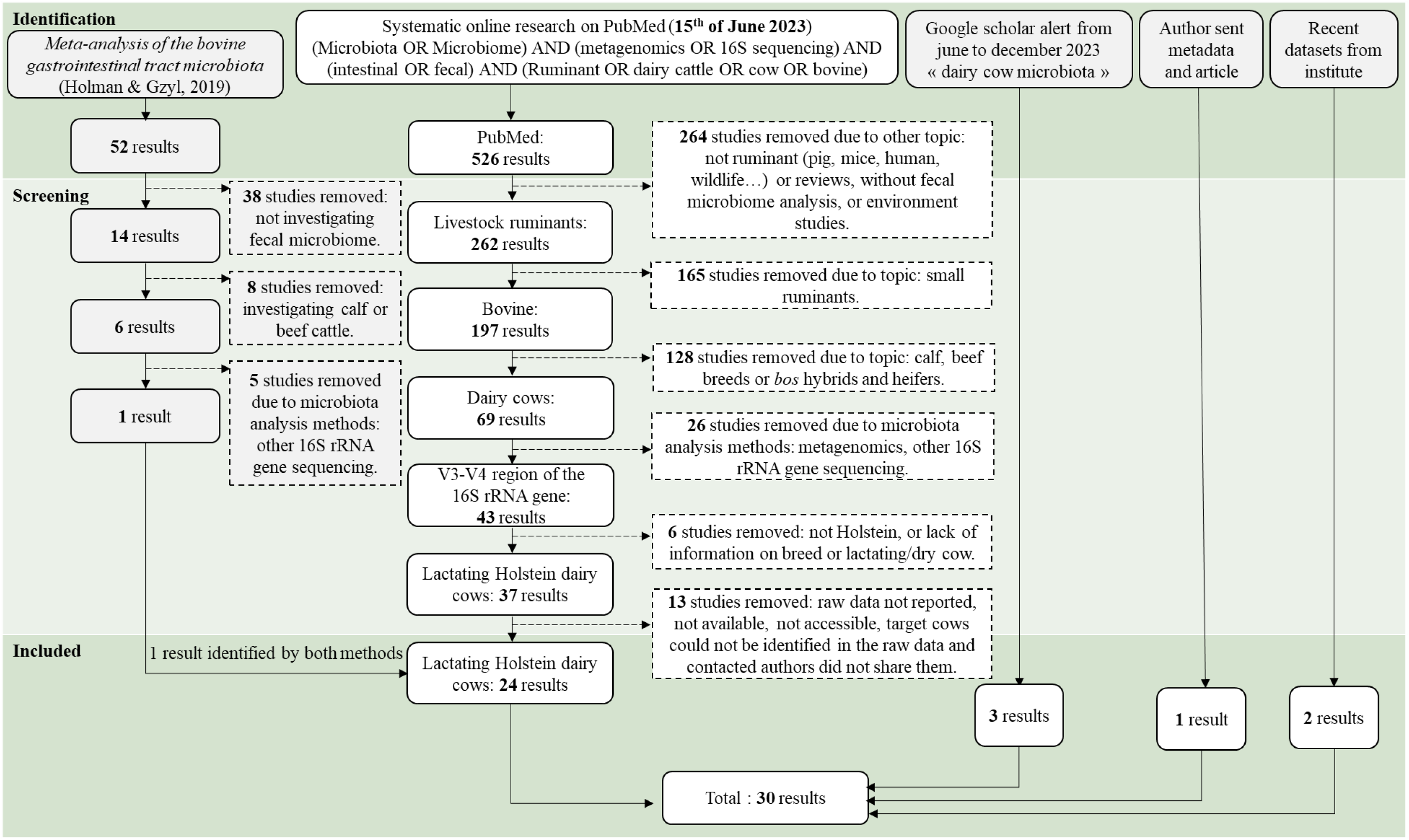
Preferred Reporting Items for Systematic Reviews and Meta-Analyses (PRISMA) flow diagram for article inclusion

The metadata retrieved was found to be inconsistent with little information on the animals available in the articles. Information regarding factors such as parity, lactation stage, ration, feed intake, or health status was indeed lacking for some cows. This prevented a thorough description of the entire dataset and differential analysis according to age, lactation rank or health status.

For the recovered data, different bioinformatics processing methods were applied; the process and results are indicated in **Supplementary Tables 1 and 2**.

The characteristics of the 34 *phyloseq* objects produced are presented in **Table 2**.

**Table 2:** Characteristics of the output of the different bio-informatic processes

|  | Cluster number | Sample number |
| --- | --- | --- |
| (SEYAMA ET AL., 2016) | 1,307 | 4 |
| (SUZUKI ET AL., 2023) | 1,290 | 38 |
| (HUANG, JI, WANG, ET AL., 2020)_27SAM | 1,329 | 27 |
| (HUANG, JI, WANG, ET AL., 2020)_S7-S8-S9 | 1,299 | 3 |
| (DANKWA ET AL., 2021) | 1,890 | 28 |
| (WANG, ZHAO, ET AL., 2022) | 1,482 | 23 |
| (GAOWA ET AL., 2021) | 1,973 | 11 |
| (SCARSELLA ET AL., 2021)-50SAMPLES | 1,662 | 50 |
| (SCARSELLA ET AL., 2021)-S53-S51-S36-S27-S49 | 1,485 | 5 |
| (WANG, NAN, ET AL., 2022A)-10SAMPLES | 1,920 | 10 |
| (WANG, NAN, ET AL., 2022A)-20SAMPLES | 1,886 | 20 |
| (YANG ET AL., 2022) | 1,139 | 38 |
| (TARDÓN ET AL., 2022) | 1,336 | 20 |
| (LEE ET AL., 2023) | 1,553 | 18 |
| (F. ZHANG ET AL., 2022) | 2,567 | 24 |
| (Z. LUO ET AL., 2022) | 1,788 | 20 |
| (LYU ET AL., 2022) | 2,299 | 14 |
| (SLANZON ET AL., 2022) | 1,296 | 209 |
| (FLORIDIA ET AL., 2023) | 1,172 | 16 |
| (JIANG ET AL., 2023) | 1,665 | 12 |
| (HAO ET AL., 2023) | 2,303 | 24 |
| (LI ET AL., 2023) | 255 | 29 |
| (OSSA-TRUJILLO ET AL., 2023) | 886 | 64 |
| (SUN ET AL., 2017) | 761 | 30 |
| (NOEL ET AL., 2019) | 1,833 | 10 |
| (HUANG, JI, YAN, ET AL., 2020) | 924 | 5 |
| (ZHONG ET AL., 2020) | 1,567 | 49 |
| (G. ZHANG ET AL., 2019) | 1,821 | 180 |
| (DONG ET AL., 2022) | 1,856 | 8 |
| (J. ZHANG ET AL., 2022)-4SAMPLES | 282 | 4 |
| (J. ZHANG ET AL., 2022)-R1-14SAMPLES | 982 | 14 |
| (WANG, NAN, ET AL., 2022B) | 1,261 | 45 |
| (BRULIN ET AL., 2024) | 1,985 | 697 |
| (ARNALOT ET AL., *ANIMAL MICROBIOME*, ACCEPTED FOR PUBLICATION) | 2,227 | 411 |

Merging the 34 *phyloseq* datasets resulted in 514 clusters at the genus level and 2,160 samples. *Archaea* (n=5), unclassified kingdom (n=1) and undefined bacterial phyla (n=2) were discarded. A total of 24 samples yielding fewer than 1,000 sequences were removed; including those from Brulin et al., (2024): ERR13304407, ERR13304726, ERR13304740,ERR13304742, from Dankwa et al., (2021): SRR12183083, SRR12183104, SRR12183106, SRR12183108, SRR12183110, from Li et al., (2023): SRR26046271, SRR26046272, SRR26046273, SRR26046277, SRR26046279, SRR26046290, SRR26046296, SRR26046297, SRR26046300, SRR26046301, SRR26046302 and from J. Zhang et al., (2022): SRR9901338, SRR9901342, SRR9901344, SRR9901345.

The final *phyloseq* dataset comprise 506 clusters and 2,136 samples.

The number of sequences and clusters per sample varied considerably between studies. In addition to this disparity, it is important to note that, during the bioinformatic processing, certain samples did not share any sequences with others from the same study, despite likely being processed in the same way (DNA extraction, amplification, and sequencing run).

After implementing the denoising process, the genus-level dataset, which initially contained 506 clusters, produced 185 clusters for the 2,136 samples at the genus level. The number of sequences per sample ranged from 1,060 to 965,946, with a mean of 42,986. After filtering, the range was 229 to 963,739, with a mean of 42,783 sequences. The cluster size (number of sequences per genus) originally ranged from 19 to 12,763,111, with a mean of 181,459. After filtering, this ranged from 9,428 to 12,763,111 sequences per genus, with a mean of 493,974.

### Fecal microbiota diversity and composition

The observed richness exhibited significant variation between articles (**Supplementary Figure 1**). This makes it difficult to identify discernible trends across the diverse sample sets. A similar outcome was observed concerning beta diversity (**Supplementary Figure 2**).

A total of 11 bacterial phyla were detected, with the *Firmicutes* phylum identified as the most abundant, accounting for an average of 66.4% of the total bacteria, followed by the *Bacteroidota* phylum, accounting for 21.9%. Other notable phyla included *Proteobacteria* (5.00%,), *Actinobacteriota* (3.77%) and *Spirochaetota* (1.56%).

Furthermore, several low-abundance phyla were consistently detected in over 1,000 samples, including *Patescibacteria* (0.67%), *Verrucomicrobiota* (0.41%) and *Cyanobacteria* (0.10%). Other notable phyla included *Fibrobacterota* (0.07%), *Desulfobacterota* (0.06%) and *Planctomycetota* (0.03%) were detected in fewer than half of the 2,136 analyzed samples (958 and 563 samples, respectively).

### Identification of the core microbiota in lactating dairy cattle

The core microbiota was investigated at the family level (**Figure 2**).

**Figure 2:**
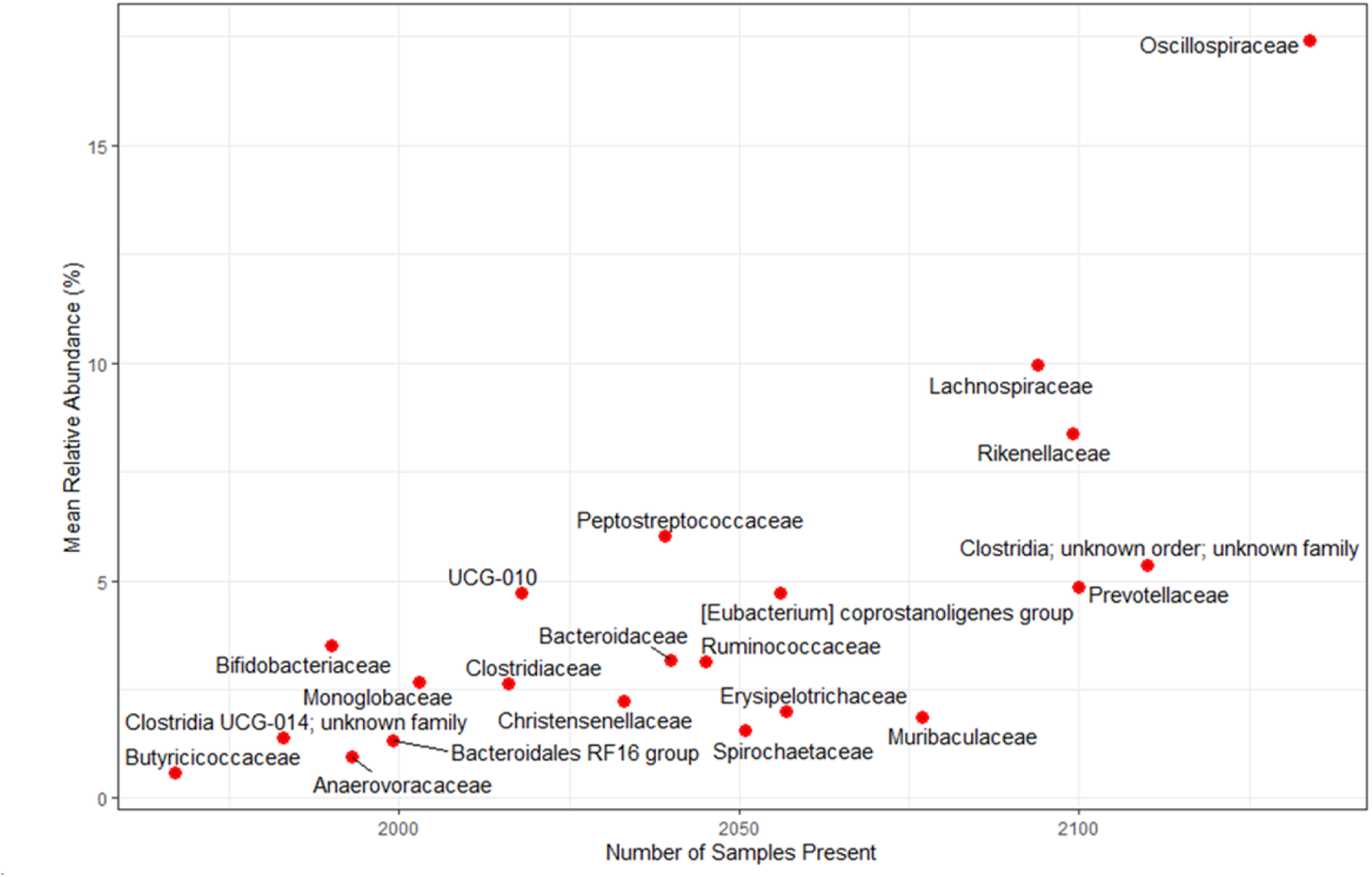
Core microbiota plot for the 2,136 samples across the 81 families (21 of which were identified as part of the core microbiota shown on the prevalence versus abundance plot).

The families *Oscillospiraceae*, *Prevotellaceae, Rikenellaceae*, *Lachnospiraceae*, *Muribaculaceae, Erysipelotrichaceae, [Eubacterium] coprostanoligenes* group, *Spirochaetaceae*, *Ruminococcaceae*, *Bacteroidaceae, Peptostreptococcaceae*, and *Christensenellaceae, UCG-010* from the *Oscillospirales* order, *Clostridiaceae*, *Monoglobaceae*, *Bacteroidales RF16* group, *Anaerovoraceae*, *Bifidobacteriaceae*, *Clostridia UCG-014*; unknown family, and *Butyricicoccaceae* were identified as part of the core microbiota. Excluding the *Clostridia; unknown order; unknown family* and *Clostridia UCG-014; unknown family,* these taxa represented 23% of the identified families in the dataset and 82% of the total relative abundance.

### Identification and description of different microbiota profiles

As shown in **Figure 3**, a strong source or article effect was observed. To alleviate this effect, a PCA-based correction was applied to the dataset after performing a CLR transformation.

**Figure 3:**
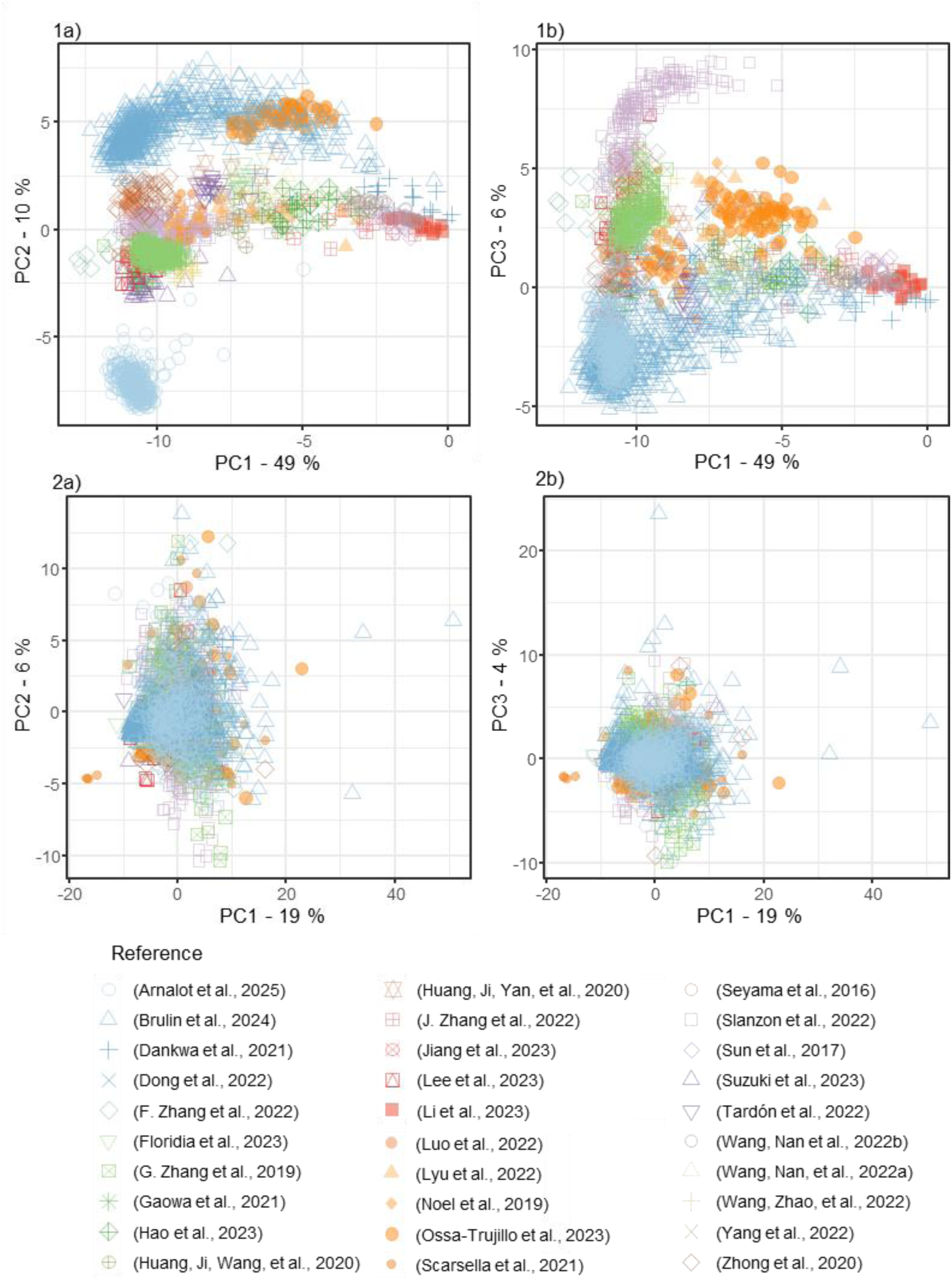
Principal component analysis (PCA) after centered log ratio (CLR) transformation not corrected (1a and 1b) and corrected with multi-group analysis according to the study (2a and 2b).

Following the correction of the article, the effect of microbiota cluster could be identified. The choice of three clusters is supported by the hierarchical clustering dendrogram using the corrected dataset (distance = 1-correlation of the first 50 components of the corrected PCA, which explains 71% of the variance, using Ward’s method with squared distances as linkage method). A natural separation emerged at this level (**Figure 4**). The distribution of the samples across the microbiota profiles in the included articles is presented in **Supplementary Table 4**.

**Figure 4:**
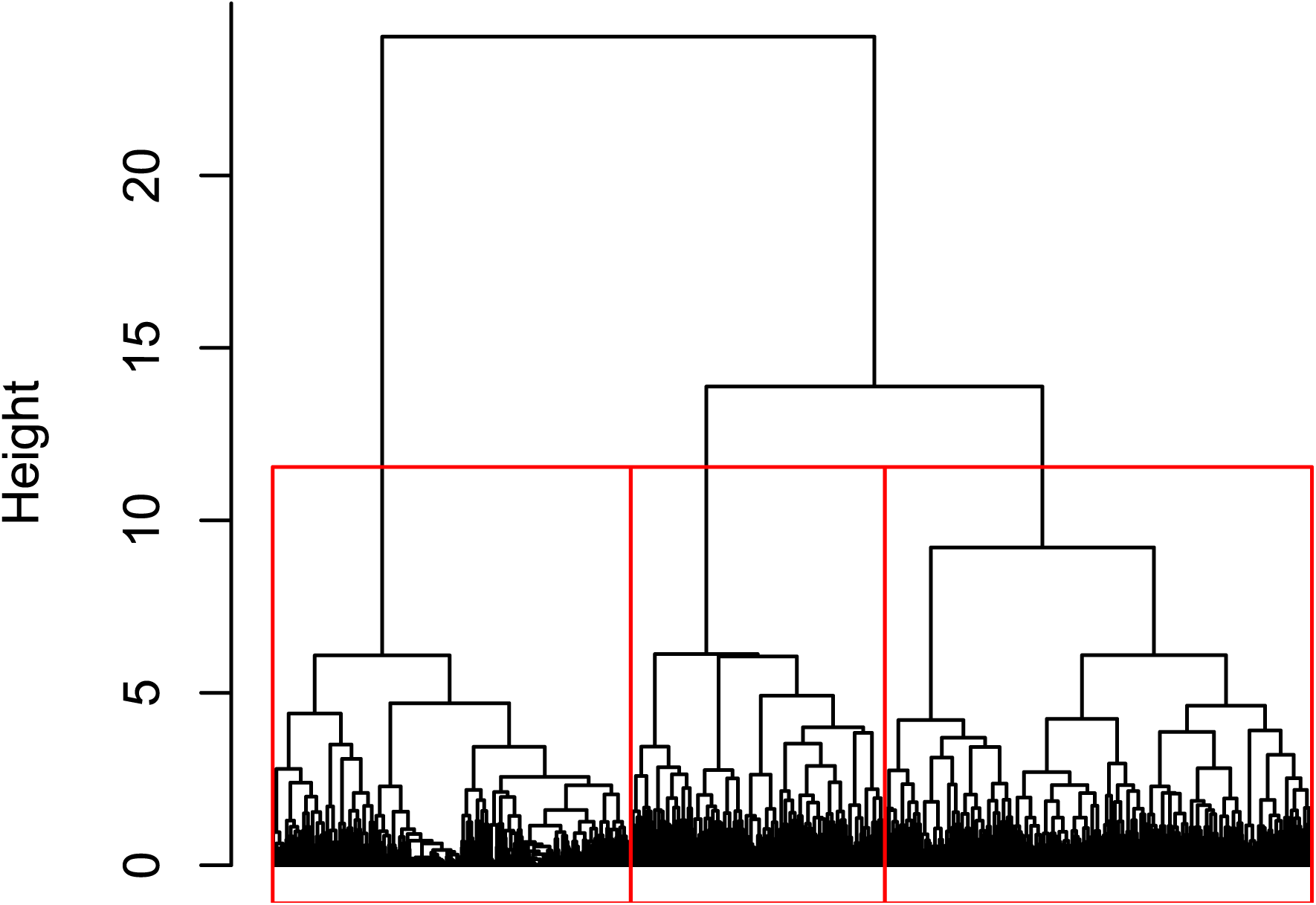
Dendrogram determining the three microbiota profiles. The distance is based on 1-correlation of the first 50 components of the corrected PCA, which explains 71% of the variance, using Ward’s linkage method with squared distances.

As shown in **Figure 5**, the ANCOM-BC analysis revealed substantial disparities among the profiles.

**Figure 5:**
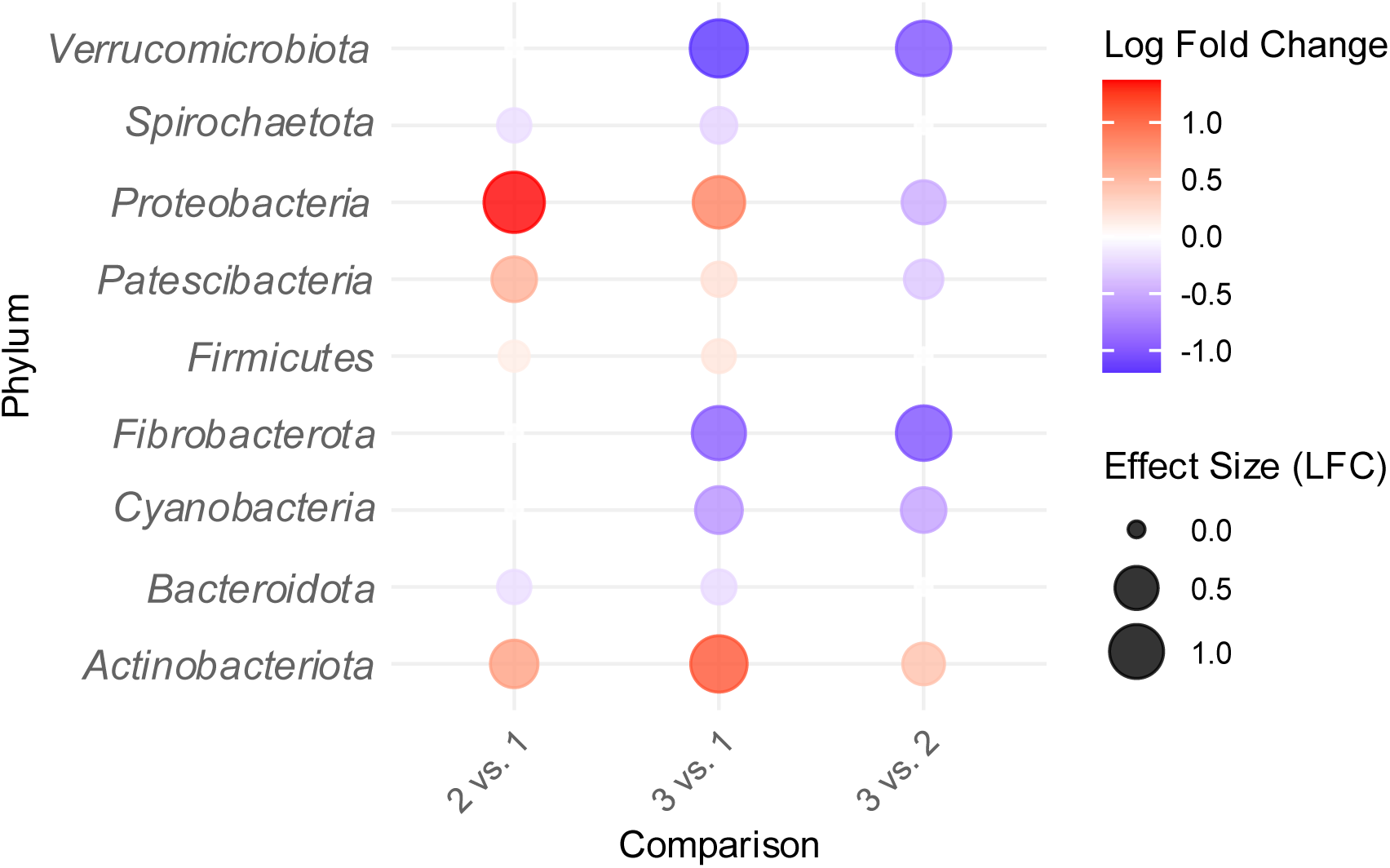
ANCOM-BC output for the pairwise comparison of the three microbiota profiles identified at the phylum level, exclusively for significant log fold change (LFC). The size of the dot corresponds to the magnitude of the LFC. Negative LFC is indicated by blue, while positive LFC is indicated by red.

**Profile 1** exhibited a decrease in the abundance of *Proteobacteria* and *Actinobacteriota*, accompanied by a modest decrease in *Patescibacteria*. In contrast, a significant increase of *Verrucomicrobiota*, *Cyanobacteria*, and *Fibrobacterota* was observed, compared to **Profile 3**. **Profile 2** exhibited a higher abundance of *Proteobacteria* compared to the other profiles; however, for other phyla, it was mainly an intermediate between the other two profiles.

**Profile 3** was characterized by elevated levels of *Actinobacteriota* andlower levels of *Verrucomicrobiota*, *Cyanobacteria*, and *Fibrobacterota*.

The two phyla with the lowest abundance, *Desulfobacterota* and *Planctomycetota,* did not exhibit any difference across microbiota profiles.

The differential abundance of bacterial species across microbial profiles was also determined at the family level (**Figure 6**). **Profile 1** was enriched in *Flavobacteriaceae*, *Fibrobacteraceae*, and *Gammaproteobacteria; unknown order; unknown family*. However, it depleted in *Bifidobacteriaceae, Clostridiaceae and Akkermansiaceae* compared to **Profile 3**.

**Figure 6:**
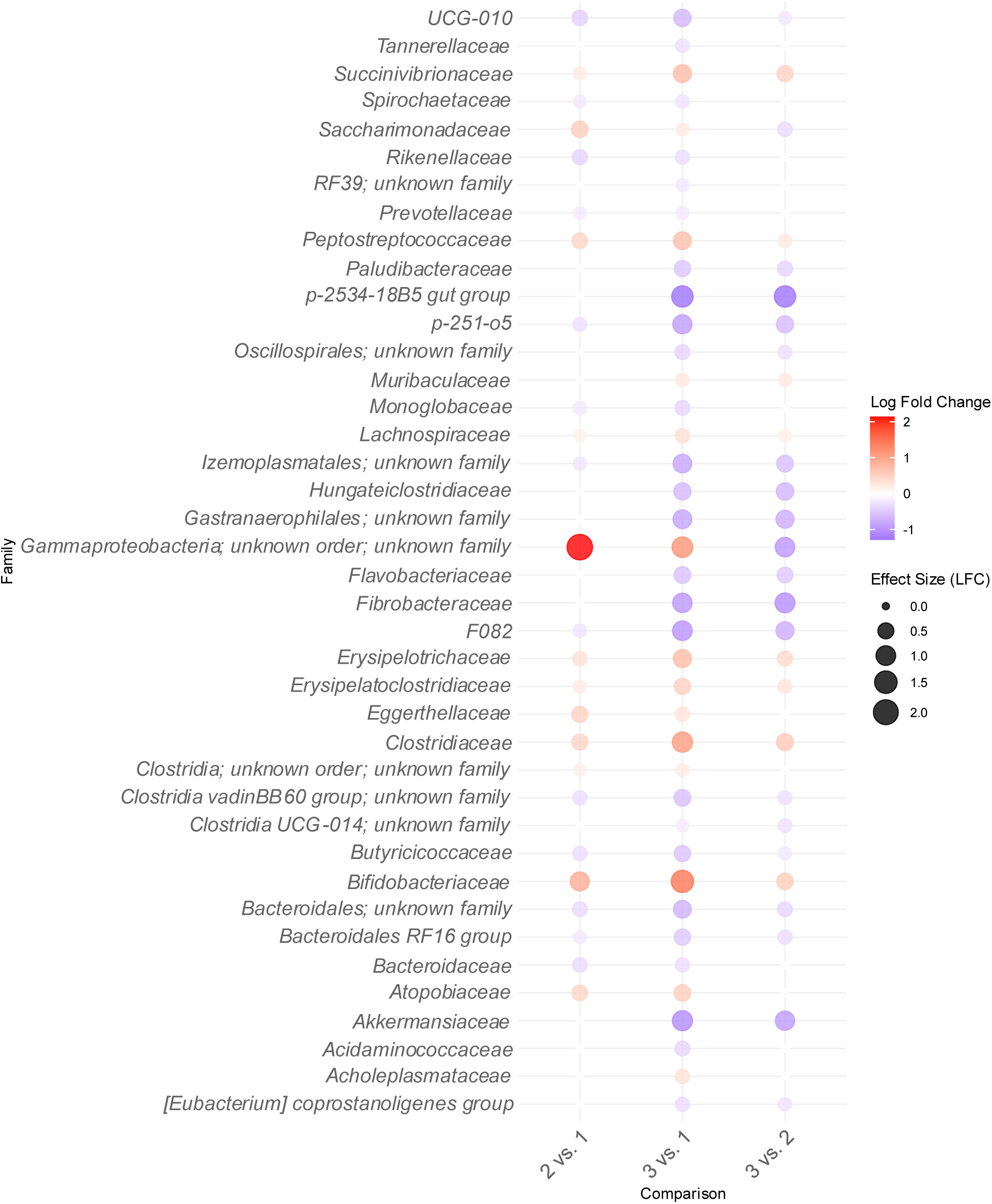
ANCOM-BC output for pairwise comparison of the three microbiota profiles identified at the family level, exclusively for significant log fold change (LFC). The size of the dot corresponds to the magnitude of the LFC. Negative LFC is indicated by blue, while positive LFC is indicated by red.

**Profile 2** seemed to be more of a transitional profile between the others.

## DISCUSSION

This study investigated the fecal microbiota of lactating Holstein dairy cows by examining existing literature. Following corrections of the variability due to data sources, three different microbiota profiles were identified according to these data, as well as 21 microbiota families that constitute the overall core microbiota.

Among the recovered data from publication search and direct contact with the authors, we had to apply different bioinformatic processing methods for several articles, as the reported primers were incompatible with the analytical tool. Additionally, some samples had to be excluded due to inconsistencies within the same article, such as atypical read lengths and discrepancies between read1 and read2. These issues suggest that the quality of certain samples may be questionable. The variability in the quality of the data can also be discussed as the number of sequences per sample and shared sequences between samples from the same origin varied considerably across studies.

To minimize the variability previously reported by Holman et al. (2017) and Holman and Gzyl. (2019), the investigation was conducted exclusively on the same hypervariable region. Nevertheless, there was a strong variability across samples according to the source of origin, which led us to use batch correction *via* multivariate integration, a process combining multiple independent studies that assessed the same predictors (Rohart, Eslami, et al., 2017; Rohart, Gautier, et al., 2017).

**Profile 1** was characterized by a significant depletion of *Actinobacteriota,* including the family *Bifidobacteriaceae*, known to have beneficial functions in carbohydrate fermentation (Turroni et al., 2011), suggesting a potential association of this profile with negative outcomes. Furthermore, the slightly reduced levels of *Firmicutes*, along with the associated *Lachnospiraceae*, indicated a limited capacity for fiber degradation compared to other profiles (Flint et al., 2008).

Compared to the other profiles, **Profile 2** exhibited an increase in *Proteobacteria* and *Patescibacteria*. The main difference at the family level was related to the *Gammaproteobacteria*; unknown order; unknown family. Profile 2 appeared to be a more transient profile that is less distinct than **Profiles 1** and **3**, from each other.

**Profile 3** exhibited a preponderance of *Actinobacteriota*, specifically *Bifidobacteriaceae.* These taxa play a role in complex carbohydrate metabolism, short-chain fatty acid (SCFA) production and bile acid metabolism (Clavel et al., 2014; Turroni et al., 2011). *Firmicutes*, including *Lachnospiraceae*, also demonstrated moderate abundance, providing fibre degradation and butyrate production, which are crucial for colonocyte health and anti-inflammatory effects (Flint et al., 2008). **Profile 3** has also a lower abundance in *Akkermansiaceae* (*Verrucomicrobiota*) compared to other groups, suggesting decreased mucin degradation, which plays a key role in maintaining gut barrier integrity and modulating immune responses (Everard et al., 2013). However, *Akkermansia* has also been associated with diseased cows such as left displacement of the abomasum and subclinical mastitis (S. Luo et al., 2022; Zhao et al., 2023). A recent meta-analysis in human microbiota revealed the fact that its composition and diversity differ between world regions. The most important technical factor being related to DNA extraction and primers (Abdill et al., 2025). In our study, the majority of the samples originated from the same countries, consequently, the potential for variation due to regional differences was not assessed. However, it should be noted that the effects of diet and region may be less variable overall in dairy cows, as they are managed similarly, compared to meat cattle, where management is much more different (feedlots compared to free pasture). Unlike humans, for whom considerable variability is observed across a range of factors, including country of residence, ethnicity, lifestyle, and age, such variability is less pronounced in dairy cows.

Based on Shade and Stopnisek, (2019) description, Neu, Allen and Roy, (2021) hypothesized that investigating the core microbiota at a higher taxonomic level might be more relevant than at a lower one (e.g., species level). We identified several families as part of the core microbiota of the lactating Holstein dairy cow. Interestingly, the core constituted merely a minority fraction of the families(representing only 23% of the entire families in the dataset), yet it accounts for the majority of the relative abundance of the samples, leaving only 18% of the relative abundance to other taxa.

We would also like to emphasize the challenges involved in conducting this meta-analysis. In the current era of Open Science — defined as an effort to make scientific research more accessible, transparent, reproducible, and reliable (Bertram et al., 2023) — we encountered persistent issues with data availability. In numerous cases, the raw data were not publicly accessible, essential metadata were missing, or the authors did not respond to data requests. As a consequence, 13 studies meeting our inclusion criteria could not be incorporated into the analysis, and even the included datasets lacked complete metadata. This highlights a significant gap between the principles of Open Science and the practical reality of data sharing in microbiota research. Similar concerns have recently been raised in ecology, calling for clearer and more enforceable guidelines on data availability and management (Koivisto & Mäntylä, 2024).

The present study provides a valuable foundation for future research on the dairy cow microbiota. However, the limited availability and precision of metadata prevented a more in-depth investigation into potential associations with health status, lactation rank or lactation stage. We recommend that future studies address this limitation by ensuring that metadata are both comprehensive and publicly available, in order to facilitate more robust and generalizable conclusions on these important topics.

## CONCLUSION

This meta-analysis demonstrated the significant impact of the sample reference origin on results, highlighting the challenge of deriving global conclusions. Nevertheless, the study successfully diminished the article effect and identified three distinct microbiota profiles in lactating Holstein dairy cows. The core microbiota identified within the datasets was characterized, with 21 microbiota families representing 82% of the relative abundance.

The study also underscores the scarcity of high-quality data, in terms of both raw sequencing reads and associated metadata. While some authors shared their datasets upon request, broader data availability—including comprehensive metadata—would greatly enhance the comparability of results across studies. Such improvements are essential to generate more robust conclusions and to advance our understanding of key factors influencing cattle productivity, health and welfare.

## DATA AVAILABILITY

All data included in this article were obtained from public repositories with the following <colcnt=3> accession numbers: PRJNA1170208, PRJNA628713, PRJNA644954, PRJNA657329, PRJNA682766, PRJNA725200, PRJNA760802, PRJNA774499, PRJNA790039, PRJNA815875, PRJNA822933, PRJNA838477, PRJNA860705, PRJNA881400, PRJNA928233, PRJNA943133, PRJNA973208, PRJNA824686, PRJNA625290, PRJNA351736, PRJNA525989, PRJNA540088, PRJNA526913, PRJNA550022, PRJNA558571, PRJNA828386, PRJEB75421 from the National Centre for Biotechnology Information Sequence Red Archive and DRA004106 and DRR401736 from DNA Data Bank of Japan. Data from (Dong et al., 2022) was directly obtained from the authors.

## Authors’ contributions

A.Z., G.F., and L.A. contributed to the design of the study. G.P. and L.A. conducted the bioinformatic processing. L.A. performed the statistical analysis for which S.D. provided direction. A.Z., G.F. and L.A. interpreted the data. The first version of the manuscript was drafted by L. A., with substantial revisions thereafter by A. Z. and G. F.. All authors approved the final version submitted for publication.

## ACKNOWLEDGMENTS

Expressions of profound gratitude are extended to all authors who graciously responded to our requests and provided us with their data, thereby demonstrating remarkable receptiveness and transparency.

Additionally, we would like to thank Dr Sylvie COMBES who provided guidance in the study and Dorian SENSEBY, who assisted with the initial version of the microbiota profiles.

## ABBREVIATIONS

CLR: Centered Log Ratio
DDBJ: DNA Data Bank of Japan
ENA: European Nucleotide Archive
LFC: Log Fold Change
NCBI: National Center for Biotechnology Information
NMDS: Non-Metric Multidimensional Scaling
PCA: Principal Component Analysis
PRISMA: Preferred Reporting Items for Systematic Reviews and Meta-Analyses

## Supplementary data list

**Supplementary Table 1:** FROGS 4.1.0 parameters used for the samples treated with only read1 for 16S rRNA gene V3V4 amplicons from literature

| Tool | preprocessing | clustering | remove_chimera | cluster_filters | taxonomic_affiliation |
| --- | --- | --- | --- | --- | --- |
| Parameter for R1 of the 16S rRNA V3V4 | illumina<br>--already-contiged<br>--without-primers<br>--min-amplicon-size 150<br>--max-amplicon-size 400 | --fastidious --distance 1 | default | --min-abundance 0.00005<br>--contaminant phiX | --rdp<br>--reference silva_138.1_16S.fasta |
| *(Seyama et al., 2016) - 4 samples* | 576,846 raw sequences<br>576,468 pre-processed sequences (99.9% retained)<br>number of read from 66,404 to 191,208 | total of 280,053 clusters and 576,468 sequences retained | total of 276,588 clusters and 571,762 sequences retained | total of 1,307 clusters and 275,255 sequences retained | 100% clusters and sequences affiliated |
| *(Dankwa et al., 2021) - 34 samples* | 1,593,808 raw sequences<br>1,593,808 pre-processed sequences (100.0% retained)<br>number of read from 4,577 to 248,310 | total of 648,316 clusters and 1,593,808 sequences retained | total of 563,103 clusters and 1,483,960 sequences retained | total of 1,890 clusters and 739,088 sequences retained | 100% clusters and sequences affiliated |
| *(Scarsella et al., 2021) - 5 samples* | 2,840,016 raw sequences<br>1,728,169 pre-processed sequences (60.9 % retained)<br>number of read from 15,438 to 1,061,194 | total of 393,759 clusters and 1,728,169 sequences retained | total of 338,120 clusters and 1,661,803 sequences retained | total of 1,485 clusters and 1,242,579 sequences retained | One cluster without affiliation (99.9% affiliated) and 679 sequences without affiliation (99.9% affiliated) |
| *(Wang, Nan, et al., 2022a)- 20 samples* | 833,006 raw sequences<br>833,006 pre-processed sequences (100.0 % retained)<br>number of read from 32,607 to 49,881 | total of 410,797 clusters and 833,006 sequences retained | total of 237,468 clusters and 650,291 sequences retained | total of 1,886 clusters and 269,269 sequences retained | 100% clusters and sequences affiliated |
| *(Lyu et al., 2022)- 14 samples* | 687,265 raw sequences<br>668,802 pre-processed sequences (97.3 % retained)<br>number of read from 37,002 to 60,119 | total of 256,574 clusters and 668,802 sequences retained | total of 136,943 clusters and 535,781 sequences retained | total of 2,299 clusters and 300,135 sequences retained | 100% clusters and sequences affiliated |
| *(Slanzon et al., 2022)- 209 samples* | 11,315,083 raw sequences<br>11,206,074 pre-processed sequences (99.0 % retained)number of read from 35,504 to 74,390 | total of 4,459,565 clusters and 11,206,074 sequences retained | total of 3,417,622 clusters and 9,900,184 sequences retained | total of 1,296 clusters and 5,615,151 sequences retained | 100% clusters and sequences affiliated |
| *(Hao et al., 2023)- 24 samples* | 1,566,496 raw sequences<br>1,566,417 pre-processed sequences (100.0 % retained)<br>number of read from 36,373 to 73,441 | total of 585,952 clusters and 1,566,417 sequences retained | total of 316,397 clusters and 1,250,604 sequences retained | total of 2,303 clusters and 664,641 sequences retained | 100% clusters and sequences affiliated |
| *(Ossa-Trujillo et al., 2023)- 64 samples* | 8,732,195 raw sequences<br>8,712,094 pre-processed sequences (99.8 % retained)<br>number of read from 21,865 to 246,931 | total of 2,726,371 clusters and 8,712,094 sequences retained | total of 2,539,469 clusters and 8,398,236 sequences retained | total of 886 clusters and 5,398,440 sequences retained | 100% clusters and sequences affiliated |
| *(Noel et al., 2019) - 10 samples* | 380,517 raw sequences<br>379,356 pre-processed sequences (99.7 % retained)<br>number of read from 23,283 to 65,256 | total of 99,588 clusters and 379,356 sequences retained | total of 73,663 clusters and 351,937 sequences retained | total of 1,833 clusters and 260,116 sequences retained | 15 clusters without affiliation (99.2% affiliated) and 427 sequences without affiliation (99.8% affiliated) |
| *(Huang, Ji, Yan, et al., 2020)- 5 samples* | 282,332 raw sequences<br>234,550 pre-processed sequences (83.1 % retained)<br>number of read from 43,521 to 49,336 | total of 126,014 clusters and 234,550 sequences retained | total of 74,679 clusters and 172,637 sequences retained | total of 924 clusters and 95,248 sequences retained | 100% clusters and sequences affiliated |
| *(Zhong et al., 2020)- 49 samples* | 2,728,486 raw sequences<br>2,728,486 pre-processed sequences (100.0 % retained)<br>number of read from 32,218 to 78,933 | total of 1,239,914 clusters and 2,728,486 sequences retained | total of 503,867 clusters and 1,896,006 sequences retained | total of 1,567 clusters and 1,238,694 sequences retained | 100% clusters and sequences affiliated |
| *(J. Zhang et al., 2022)- 14 samples* | 654,960 raw sequences<br>654,867 pre-processed sequences (100.0 % retained)<br>number of read from 33,757 to 61,516 | total of 424,582 clusters and 654,867 sequences retained | total of 382,831 clusters and 600,226 sequences retained | total of 982 clusters and 132,359 sequences retained | 100% clusters and sequences affiliated |

**Supplementary Table 2:**
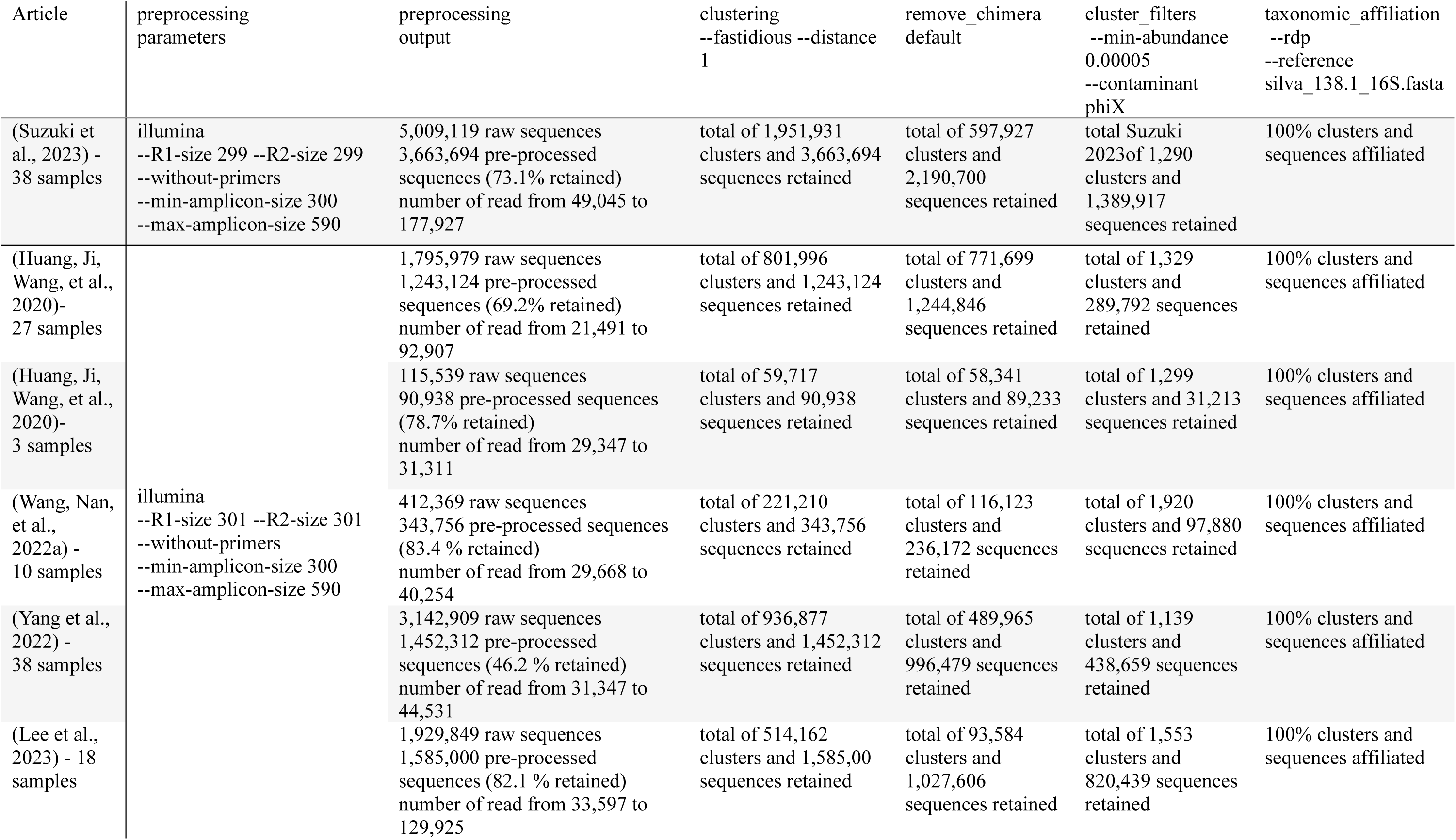
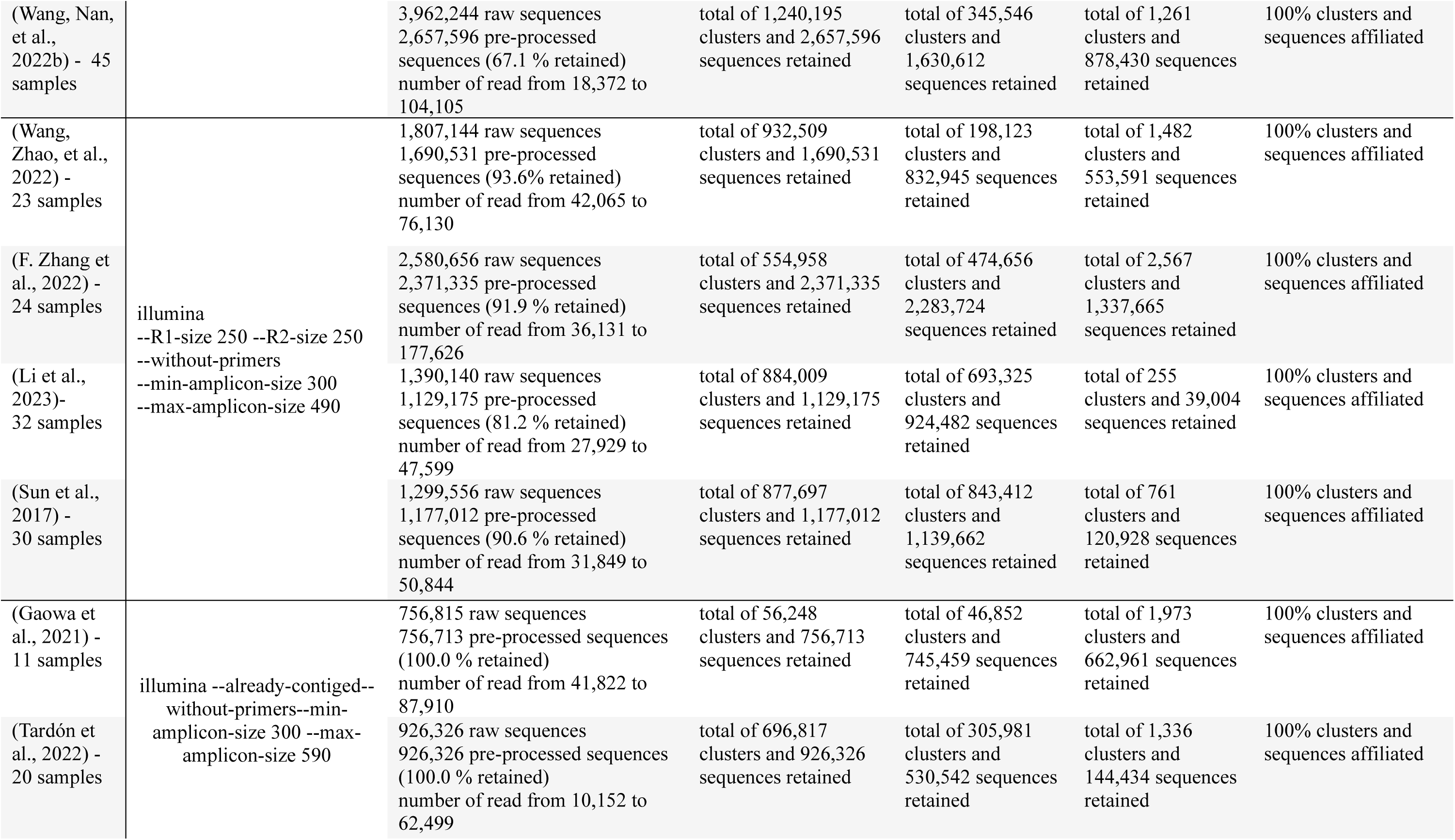
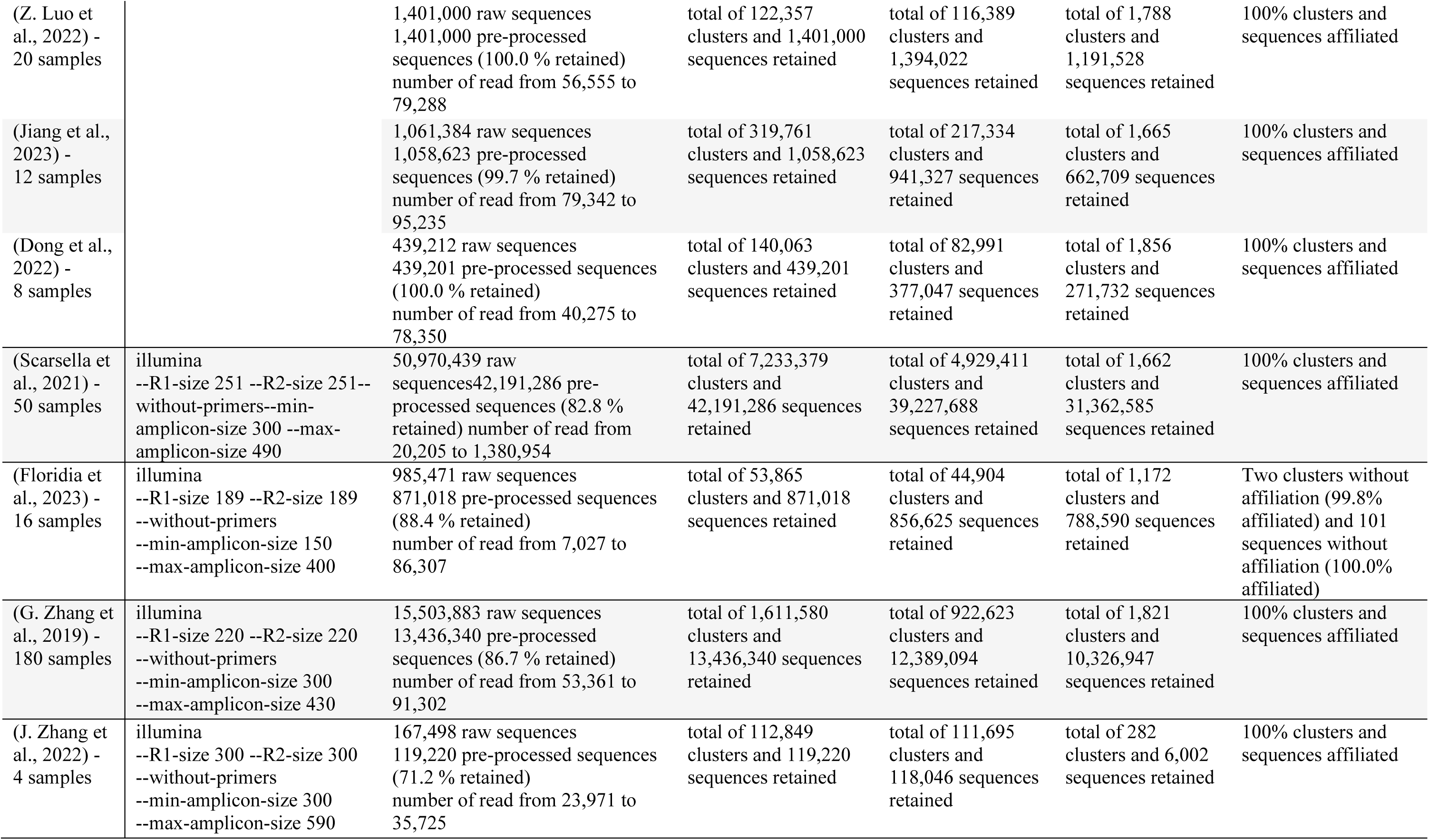
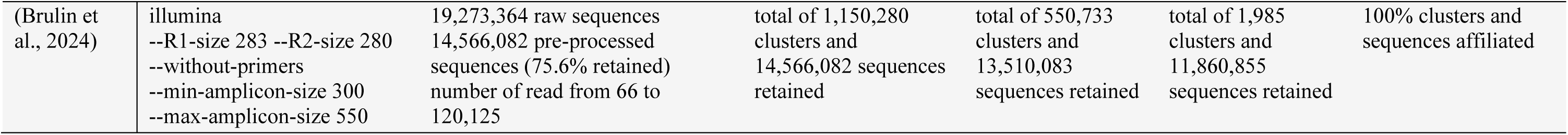
FROGS 4.1.0 parameters used for the samples treated with both reads for 16S rRNA gene V3V4 amplicons from literature

**Supplementary Table 3:**
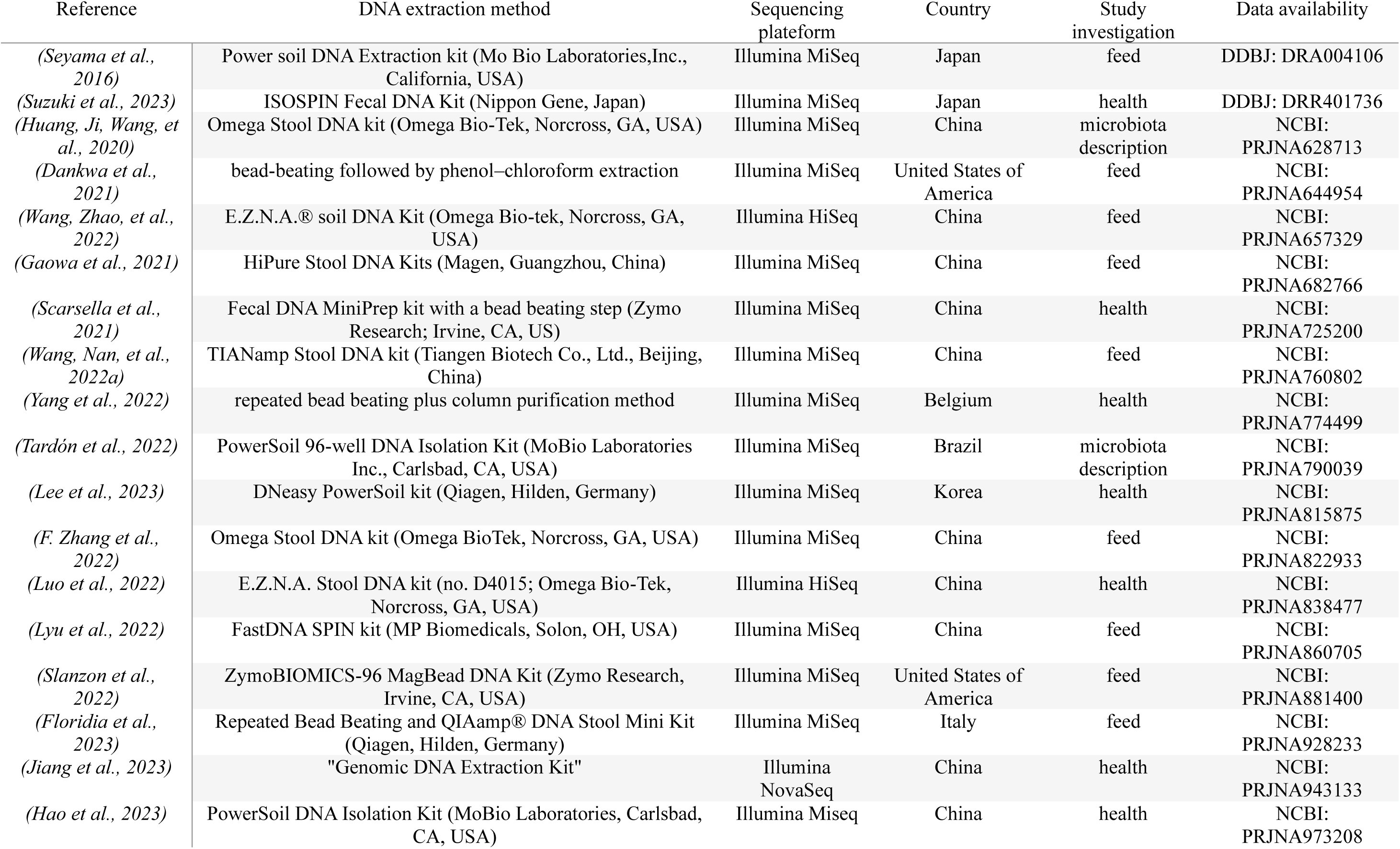
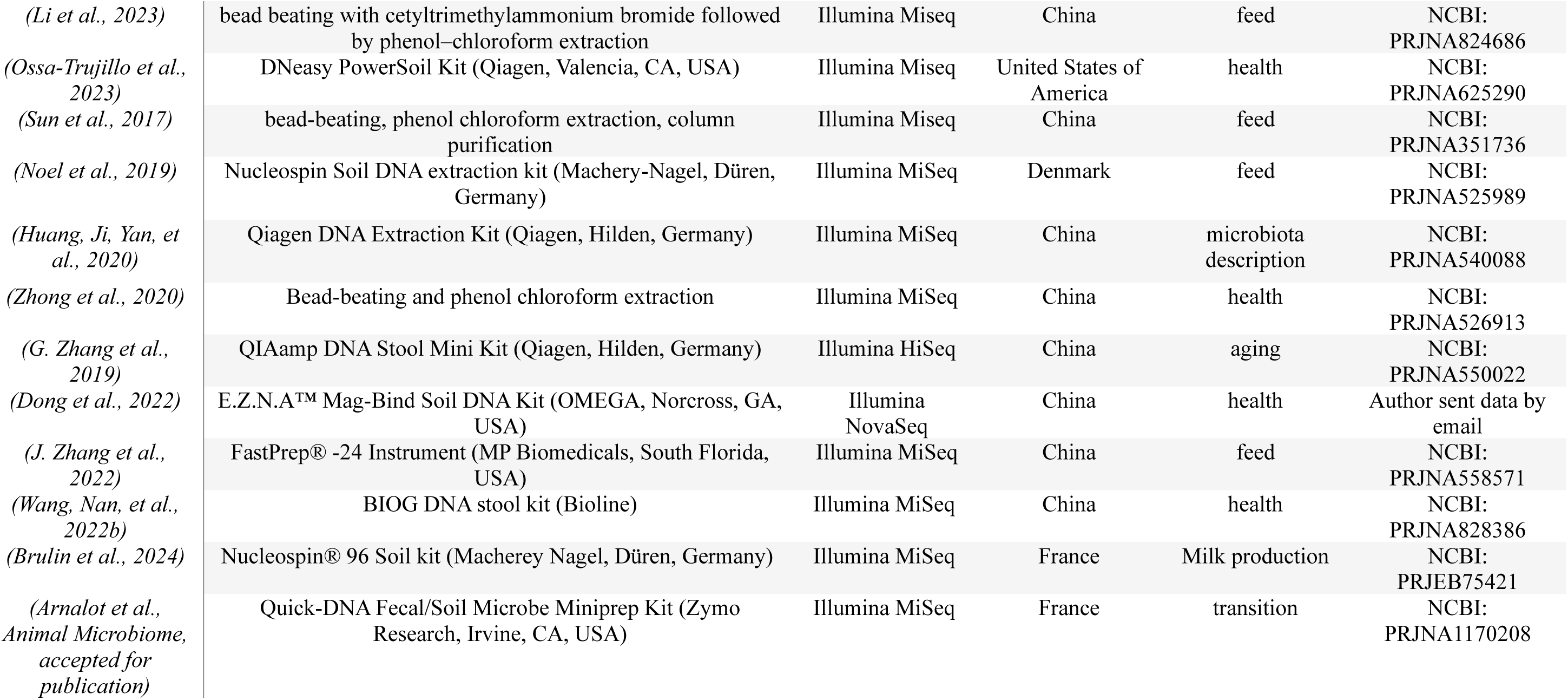
Information on the studies included in the meta-analysis

**Supplementary Table 4:** Number of samples per microbiota profile according to the reference of origin

| Microbiota Profiles | 1 | 2 | 3 |
| --- | --- | --- | --- |
| (Arnalot et al., Animal Microbiome, accepted for publication) | 148 | 181 | 82 |
| (Brulin et al., 2024) | 292 | 326 | 75 |
| (Dankwa et al., 2021) | 6 | 17 | 0 |
| (Dong et al., 2022) | 3 | 3 | 2 |
| (F. Zhang et al., 2022) | 7 | 13 | 4 |
| (Floridia et al., 2023) | 8 | 7 | 1 |
| (G. Zhang et al., 2019) | 66 | 69 | 45 |
| (Gaowa et al., 2021) | 4 | 5 | 2 |
| (Hao et al., 2023) | 8 | 12 | 4 |
| (Huang, Ji, Wang, et al., 2020) | 12 | 15 | 3 |
| (Huang, Ji, Yan, et al., 2020) | 3 | 2 | 0 |
| (J. Zhang et al., 2022) | 5 | 7 | 2 |
| (Jiang et al., 2023) | 4 | 3 | 5 |
| (Lee et al., 2023) | 8 | 7 | 3 |
| (Li et al., 2023) | 5 | 9 | 4 |
| (Luo et al., 2022) | 8 | 8 | 4 |
| (Lyu et al., 2022) | 5 | 5 | 4 |
| (Noel et al., 2019) | 3 | 6 | 1 |
| (Ossa-Trujillo et al., 2023) | 15 | 32 | 17 |
| (Scarsella et al., 2021) | 24 | 23 | 8 |
| (Seyama et al., 2016) | 2 | 1 | 1 |
| (Slanzon et al., 2022) | 87 | 77 | 45 |
| (Sun et al., 2017) | 8 | 15 | 7 |
| (Suzuki et al., 2023) | 16 | 16 | 6 |
| (Tardón et al., 2022) | 10 | 6 | 4 |
| (Wang, Nan et al., 2022b) | 10 | 21 | 14 |
| (Wang, Nan, et al., 2022a) | 10 | 15 | 5 |
| (Wang, Zhao, et al., 2022) | 7 | 8 | 8 |
| (Yang et al., 2022) | 17 | 13 | 8 |
| (Zhong et al., 2020) | 12 | 27 | 10 |
| (Zhong et al., 2020) | 12 | 27 | 10 |

**Supplementary Figure 1:**
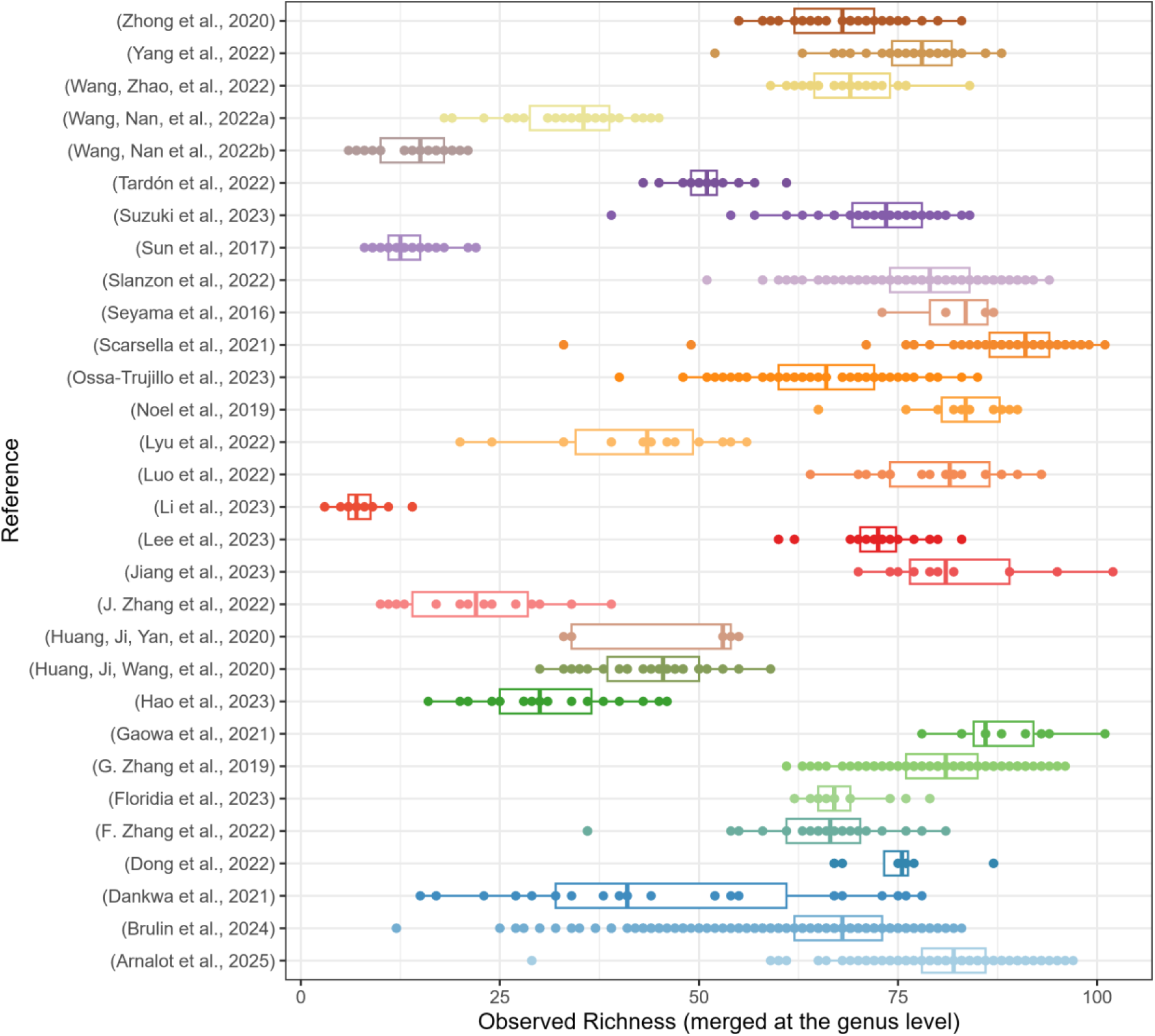
Observed richness according to the reference of origin

**Supplementary Figure 2:**
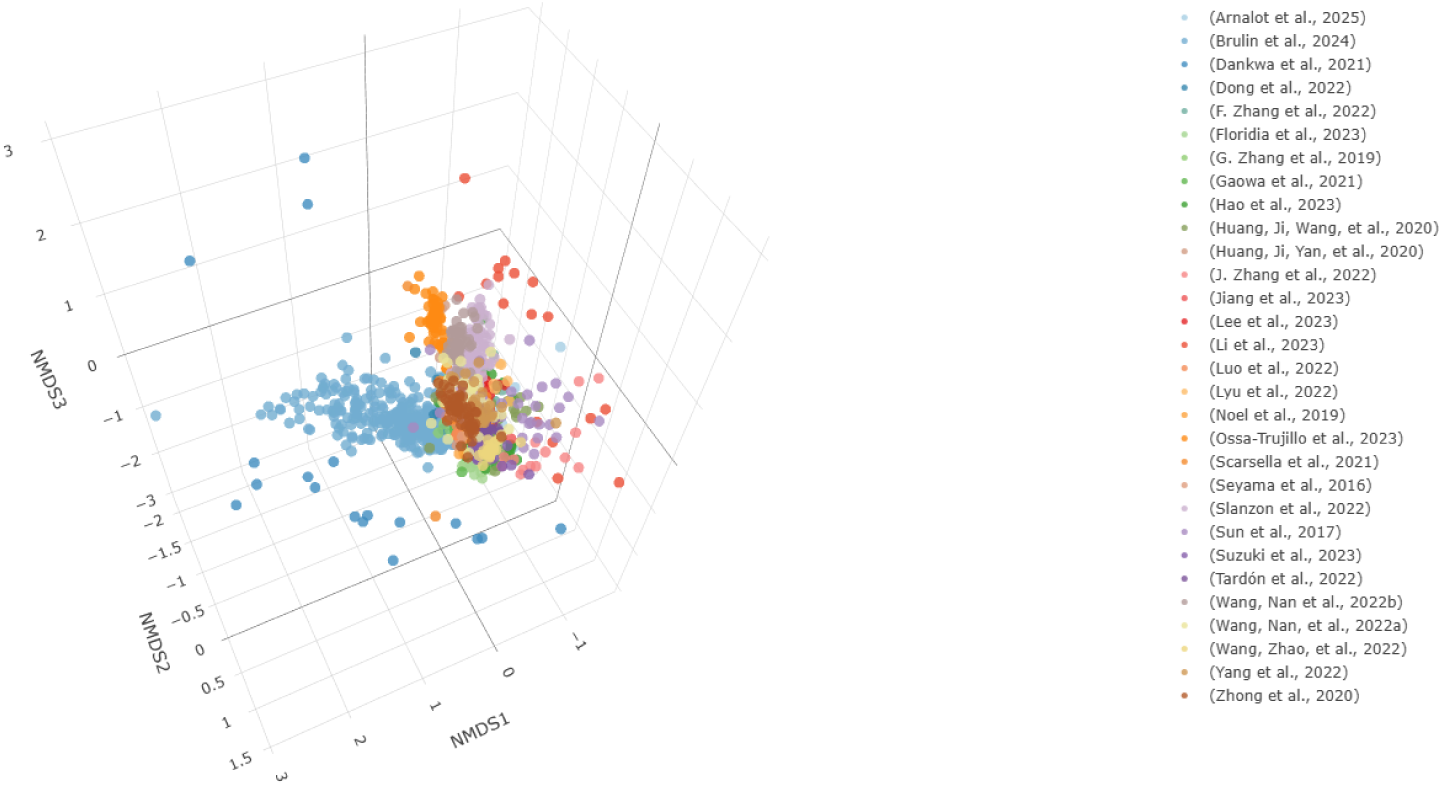
Bray-Curtis distance according to the reference of origin

## REFERENCES

1. Abdill, R. J., Graham, S. P., Rubinetti, V., Ahmadian, M., Hicks, P., Chetty, A., McDonald, D., Ferretti, P., Gibbons, E., Rossi, M., Krishnan, A., Albert, F. W., Greene, C. S., Davis, S., & Blekhman, R. (2025). Integration of 168,000 samples reveals global patterns of the human gut microbiome. Cell, 188(4), 1100–1118.e17. 10.1016/j.cell.2024.12.017

2. Bertram, M. G., Sundin, J., Roche, D. G., Sánchez-Tójar, A., Thoré, E. S. J., & Brodin, T. (2023). Open science. In Current Biology (Vol. 33, Issue 15, pp. R792–R797). Cell Press. 10.1016/j.cub.2023.05.036

3. Brulin, L., Ducrocq, S., Even, G., Sanchez, M. P., Martel, S., Merlin, S., Audebert, C., Croiseau, P., & Estellé, J. (2024). Short communication: Bifidobacterium abundance in the faecal microbiota is strongly associated with milk traits in dairy cattle. Animal, 18(8). 10.1016/j.animal.2024.101243

4. Clavel, T., Desmarchelier, C., Haller, D., Gérard, P., Rohn, S., Lepage, P., & Daniel, H. (2014). Intestinal microbiota in metabolic diseases: From bacterial community structure and functions to species of pathophysiological relevance. Gut Microbes, 5(4), 544–551. 10.4161/gmic.29331

5. Dankwa, A. S., Humagain, U., Ishaq, S. L., Yeoman, C. J., Clark, S., Beitz, D. C., & Testroet, E. D. (2021). Bacterial communities in the rumen and feces of lactating Holstein dairy cows are not affected when fed reduced-fat dried distillers’ grains with solubles. Animal, 15(7). 10.1016/j.animal.2021.100281

6. Dong, L., Meng, L., Liu, H., Wu, H., Schroyen, M., Zheng, N., & Wang, J. (2022). Effect of Cephalosporin Treatment on the Microbiota and Antibiotic Resistance Genes in Feces of Dairy Cows with Clinical Mastitis. Antibiotics, 11(1). 10.3390/antibiotics11010117

7. Escudié, F., Auer, L., Bernard, M., Mariadassou, M., Cauquil, L., Vidal, K., Maman, S., Hernandez-Raquet, G., Combes, S., & Pascal, G. (2018). FROGS: Find, Rapidly, OTUs with Galaxy Solution. Bioinformatics, 34(8), 1287–1294. 10.1093/bioinformatics/btx791

8. Everard, A., Belzer, C., Geurts, L., Ouwerkerk, J. P., Druart, C., Bindels, L. B., Guiot, Y., Derrien, M., Muccioli, G. G., Delzenne, N. M., De Vos, W. M., & Cani, P. D. (2013). Cross-talk between Akkermansia muciniphila and intestinal epithelium controls diet-induced obesity. Proceedings of the National Academy of Sciences of the United States of America, 110(22), 9066–9071. 10.1073/pnas.1219451110

9. Fan, Y., & Pedersen, O. (2021). Gut microbiota in human metabolic health and disease. In Nature Reviews Microbiology (Vol. 19, Issue 1, pp. 55–71). Nature Research. 10.1038/s41579-020-0433-9

10. Flint, H. J., Bayer, E. A., Rincon, M. T., Lamed, R., & White, B. A. (2008). Polysaccharide utilization by gut bacteria: Potential for new insights from genomic analysis. In Nature Reviews Microbiology (Vol. 6, Issue 2, pp. 121–131). 10.1038/nrmicro1817

11. Floridia, V., Russo, N., D’Alessandro, E., Lopreiato, V., Pino, A., Amato, A., Liotta, L., Caggia, C., & Randazzo, C. L. (2023). Effect of olive cake supplementation on faecal microbiota profile of Holstein and Modicana dairy cattle. Microbiological Research, 277. 10.1016/j.micres.2023.127510

12. Gaowa, N., Zhang, X., Li, H., Wang, Y., Zhang, J., Hao, Y., Cao, Z., & Li, S. (2021). Effects of rumen-protected niacin on dry matter intake, milk production, apparent total tract digestibility, and faecal bacterial community in multiparous holstein dairy cow during the postpartum period. Animals, 11(3), 1–12. 10.3390/ani11030617

13. Hao, Y., Ouyang, T., Wang, W., Wang, Y., Cao, Z., Yang, H., Guan, L. L., & Li, S. (2023). Competitive Analysis of Rumen and Hindgut Microbiota Composition and Fermentation Function in Diarrheic and Non-Diarrheic Postpartum Dairy Cows. Microorganisms, 12(1), 23. 10.3390/microorganisms12010023

14. Holman, D. B., Brunelle, B. W., Trachsel, J., & Allen, H. K. (2017). Meta-analysis To Define a Core Microbiota in the Swine Gut. MSystems, 2(3). 10.1128/msystems.00004-17

15. Holman, D. B., & Gzyl, K. E. (2019). A meta-analysis of the bovine gastrointestinal tract microbiota. FEMS Microbiology Ecology, 95(6), fiz072. 10.1093/femsec/fiz072

16. Huang, S., Ji, S., Wang, F., Huang, J., Alugongo, G. M., & Li, S. (2020). Dynamic changes of the fecal bacterial community in dairy cows during early lactation. AMB Express, 10(1). 10.1186/s13568-020-01106-3

17. Huang, S., Ji, S., Yan, H., Hao, Y., Zhang, J., Wang, Y., Cao, Z., & Li, S. (2020). The day-to-day stability of the ruminal and fecal microbiota in lactating dairy cows. MicrobiologyOpen, 9(5), e990. 10.1002/mbo3.990

18. Jiang, C., Hou, X., Gao, X., Liu, P., Guo, X., Hu, G., Li, Q., Huang, C., Li, G., Fang, W., Mai, W., Wu, C., Xu, Z., & Liu, P. (2023). The 16S rDNA high-throughput sequencing correlation analysis of milk and gut microbial communities in mastitis Holstein cows. BMC Microbiology, 23(1), 180. 10.1186/s12866-023-02925-7

19. Kim, M., & Wells, J. E. (2016). A Meta-analysis of Bacterial Diversity in the Feces of Cattle. Current Microbiology, 72(2), 145–151. 10.1007/s00284-015-0931-6

20. Koivisto, E., & Mäntylä, E. (2024). Are Open Science instructions targeted to ecologists and evolutionary biologists sufficient? A literature review of guidelines and journal data policies. Ecology and Evolution, 14(7). 10.1002/ece3.11698

21. Lahti, L., & Shetty, S. (2018). Introduction to the microbiome R package. *Preprint at* https://Microbiome.Github.Io/Tutorials.

22. Lee, S.-M., Park, H.-T., Park, S., Lee, J. H., Kim, D., Yoo, H. S., & Kim, D. (2023). A Machine Learning Approach Reveals a Microbiota Signature for Infection with Mycobacterium avium subsp. *paratuberculosis* in Cattle. Microbiology Spectrum. 10.1128/spectrum.03134-22

23. Li, Z., Fan, Y., Bai, H., Zhang, J., Mao, S., & Jin, W. (2023). Live yeast supplementation altered the bacterial community’s composition and function in rumen and hindgut and alleviated the detrimental effects of heat stress on dairy cows. Journal of Animal Science, 101, skac410. 10.1093/jas/skac410

24. Lin, H., & Peddada, S. Das. (2020). Analysis of compositions of microbiomes with bias correction. Nature Communications, 11(1), 3514. 10.1038/s41467-020-17041-7

25. Luo, S., Wang, Y., Kang, X., Liu, P., & Wang, G. (2022). Research progress on the association between mastitis and gastrointestinal microbes in dairy cows and the effect of probiotics. In Microbial Pathogenesis (Vol. 173). Academic Press. 10.1016/j.micpath.2022.105809

26. Luo, Z., Yong, K., Luo, Q., Du, Z., Ma, L., Huang, Y., Zhou, T., Yao, X., Shen, L., Yu, S., Deng, J., Ren, Z., Zhang, Y., Yan, Z., Zuo, Z., & Cao, S. (2022). Altered Fecal Microbiome and Correlations of the Metabolome with Plasma Metabolites in Dairy Cows with Left Displaced Abomasum. Microbiology Spectrum, 10(6). 10.1128/spectrum.01972-22

27. Lyu, J., Yang, Z., Wang, E., Liu, G., Wang, Y., Wang, W., & Li, S. (2022). Possibility of Using By-Products with High NDF Content to Alter the Fecal Short Chain Fatty Acid Profiles, Bacterial Community, and Digestibility of Lactating Dairy Cows. Microorganisms, 10(9). 10.3390/microorganisms10091731

28. Ma, G., Jin, W., Zhang, Y., Gai, Y., Tang, W., Guo, L., Azzaz, H. H., Ghaffari, M. H., Gu, Z., Mao, S., & Chen, Y. (2025). A Meta-Analysis of Dietary Inhibitors for Reducing Methane Emissions via Modulating Rumen Microbiota in Ruminants. Journal of Nutrition. 10.1016/j.tjnut.2024.12.011

29. Maechler, M. (2018). Cluster: cluster analysis basics and extensions. R Package Version 2.0. 7–1.

30. McMurdie, P. J., & Holmes, S. (2013). phyloseq: an R package for reproducible interactive analysis and graphics of microbiome census data. PloS One, 8(4), e61217.

31. Mishra, S. P., Wang, B., Jain, S., Ding, J., Rejeski, J., Furdui, C. M., Kitzman, D. W., Taraphder, S., Brechot, C., Kumar, A., & Yadav, H. (2023). A mechanism by which gut microbiota elevates permeability and inflammation in obese/diabetic mice and human gut. Gut, 72(10), 1848–1865. 10.1136/gutjnl-2022-327365

32. Monteiro, H. F., Figueiredo, C. C., Mion, B., Santos, J. E. P., Bisinotto, R. S., Peñagaricano, F., Ribeiro, E. S., Marinho, M. N., Zimpel, R., da Silva, A. C., Oyebade, A., Lobo, R. R., Coelho, W. M., Peixoto, P. M. G., Ugarte Marin, M. B., Umaña-Sedó, S. G., Rojas, T. D. G., Elvir-Hernandez, M., Schenkel, F. S., … Lima, F. S. (2024). An artificial intelligence approach of feature engineering and ensemble methods depicts the rumen microbiome contribution to feed efficiency in dairy cows. Animal Microbiome, 6(1). 10.1186/s42523-024-00289-5

33. Monteiro, H. F., Zhou, Z., Gomes, M. S., Peixoto, P. M. G., Bonsaglia, E. C. R., Canisso, I. F., Weimer, B. C., & Lima, F. S. (2022). Rumen and lower gut microbiomes relationship with feed efficiency and production traits throughout the lactation of Holstein dairy cows. Scientific Reports, 12(1). 10.1038/s41598-022-08761-5

34. Neu, A. T., Allen, E. E., & Roy, K. (2021). Defining and quantifying the core microbiome: Challenges and prospects. Proceedings of the National Academy of Sciences, 118(51), e2104429118. 10.1073/pnas.2104429118

35. Noel, S. J., Olijhoek, D. W., McLean, F., Løvendahl, P., Lund, P., & Højberg, O. (2019). Rumen and fecal microbial community structure of holstein and Jersey dairy cows as affected by breed, diet, and residual feed intake. Animals, 9(8). 10.3390/ani9080498

36. O’Hara, E., Neves, A. L. A., Song, Y., & Guan, L. L. (2020). The Role of the Gut Microbiome in Cattle Production and Health: Driver or Passenger? Annual Review of Animal Biosciences, 8(1), 199–220. 10.1146/annurev-animal-021419-083952

37. Ossa-Trujillo, C., Taylor, E. A., Sarwar, F., Vinasco, J., Jordan, E. R., Buitrago, J. A. G., Hagevoort, G. R., Lawhon, S. D., Piñeiro, J. M., Galloway-Peña, J., Norman, K. N., & Scott, H. M. (2023). Two-Dose Ceftiofur Treatment Increases Cephamycinase Gene Quantities and Fecal Microbiome Diversity in Dairy Cows Diagnosed with Metritis. Microorganisms, 11(11). 10.3390/microorganisms11112728

38. Page, M. J., McKenzie, J. E., Bossuyt, P. M., Boutron, I., Hoffmann, T. C., Mulrow, C. D., Shamseer, L., Tetzlaff, J. M., Akl, E. A., Brennan, S. E., Chou, R., Glanville, J., Grimshaw, J. M., Hróbjartsson, A., Lalu, M. M., Li, T., Loder, E. W., Mayo-Wilson, E., McDonald, S., … Moher, D. (2021). The PRISMA 2020 statement: An updated guideline for reporting systematic reviews. In The BMJ (Vol. 372). BMJ Publishing Group. 10.1136/bmj.n71

39. Petri, R. M., Schwaiger, T., Penner, G. B., Beauchemin, K. A., Forster, R. J., McKinnon, J. J., & McAllister, T. A. (2013). Characterization of the core rumen microbiome in cattle during transition from forage to concentrate as well as during and after an acidotic challenge. PLoS ONE, 8(12). 10.1371/journal.pone.0083424

40. Plaizier, J. C., Danscher, A. M., Azevedo, P. A., Derakhshani, H., Andersen, P. H., & Khafipour, E. (2021). A grain-based sara challenge affects the composition of epimural and mucosa-associated bacterial communities throughout the digestive tract of dairy cows. Animals, 11(6). 10.3390/ani11061658

41. Plaizier, J. C., Li, S., Tun, H. M., & Khafipour, E. (2017). Nutritional models of experimentally-induced subacute ruminal acidosis (SARA) differ in their impact on rumen and hindgut bacterial communities in dairy cows. Frontiers in Microbiology, 7(JAN). 10.3389/fmicb.2016.02128

42. Rohart, F., Eslami, A., Matigian, N., Bougeard, S., & Lê Cao, K. A. (2017). MINT: A multivariate integrative method to identify reproducible molecular signatures across independent experiments and platforms. BMC Bioinformatics, 18(1). 10.1186/s12859-017-1553-8

43. Rohart, F., Gautier, B., Singh, A., & Lê Cao, K. A. (2017). mixOmics: An R package for ‘omics feature selection and multiple data integration. PLoS Computational Biology, 13(11). 10.1371/journal.pcbi.1005752

44. Scarsella, E., Zecconi, A., Cintio, M., & Stefanon, B. (2021). Characterization of microbiome on feces, blood and milk in dairy cows with different milk leucocyte pattern. Animals, 11(5). 10.3390/ani11051463

45. Seyama, T., Hirayasu, H., Yoshida, G., Ohnuma, A., Qiu, Y., Nakajima, C., Kasai, K., & Suzuki, Y. (2016). The effects of administering lactic acid bacteria sealed in a capsule on the intestinal bacterial flora of cattle. In Japanese Journal of Veterinary Research (Vol. 64, Issue 3, pp. 197–203). Hokkaido University. 10.14943/jjvr.64.3.197

46. Seyoum, Y., Greffeuille, V., Kouadio, D. K. D., Kuong, K., Turpin, W., M’Rabt, R., Chochois, V., Fortin, S., Perignon, M., Fiorentino, M., Berger, J., Burja, K., Ponce, M. C., Chamnan, C., Wieringa, F. T., & Humblot, C. (2024). Faecal microbiota of schoolchildren is associated with nutritional status and markers of inflammation: a double-blinded cluster-randomized controlled trial using multi-micronutrient fortified rice. Nature Communications, 15(1). 10.1038/s41467-024-49093-4

47. Shade, A., & Stopnisek, N. (2019). Abundance-occupancy distributions to prioritize plant core microbiome membership. In Current Opinion in Microbiology (Vol. 49, pp. 50– 58). Elsevier Ltd. 10.1016/j.mib.2019.09.008

48. Slanzon, G., Sischo, W., & McConnel, C. (2022). Contrasting Fecal Methanogenic and Bacterial Profiles of Organic Dairy Cows Located in Northwest Washington Receiving Either a Mixed Diet of Pasture and TMR or Solely TMR. Animals, 12(20). 10.3390/ani12202771

49. Sun, J., Zeng, B., Chen, Z., Yan, S., Huang, W., Sun, B., He, Q., Chen, X., Chen, T., Jiang, Q., Xi, Q., & Zhang, Y. (2017). Characterization of faecal microbial communities of dairy cows fed diets containing ensiled Moringa oleifera fodder. Scientific Reports, 7. 10.1038/srep41403

50. Susanto, I., Rahmadani, M., Pangesti, R. T., Wiryawan, K. G., Laconi, E. B., & Jayanegara, A. (2024). Effects of various plant-based condensed tannin on rumen microbiome population: A meta-analysis. IOP Conference Series: Earth and Environmental Science, 1341(1). 10.1088/1755-1315/1341/1/012061

51. Suzuki, T., Murakami, H., Uchiyama, J., Sato, R., Takemura-Uchiyama, I., Ogata, M., Sogawa, K., Ishida, H., Atipairin, A., Matsushita, O., & Nagai, M. (2023). Exploratory study of volatile fatty acids and the rumen-and-gut microbiota of dairy cows in a single farm, with respect to subclinical infection with bovine leukemia virus. Annals of Microbiology, 73(1). 10.1186/s13213-023-01737-4

52. Tardón, D. C., Hoffmann, C., Santos, F. C. R., Decaris, N., Pinheiro, F. A., Queiroz, L. L., Hurley, D. J., & Gomes, V. (2022). Relationships among indicators of metabolism, mammary health and the microbiomes of periparturient holstein cows. Animals, 12(1). 10.3390/ani12010003

53. Turroni, F., van Sinderen, D., & Ventura, M. (2011). Genomics and ecological overview of the genus Bifidobacterium. International Journal of Food Microbiology, 149(1), 37–44. 10.1016/j.ijfoodmicro.2010.12.010

54. Wang, Y., Nan, X., Zhao, Y., Jiang, L., Wang, H., Zhang, F., Hua, D., Liu, J., Yang, L., Yao, J., & Xiong, B. (2022a). Changes in the Profile of Fecal Microbiota and Metabolites as Well as Serum Metabolites and Proteome After Dietary Inulin Supplementation in Dairy Cows With Subclinical Mastitis. Frontiers in Microbiology, 13. 10.3389/fmicb.2022.809139

55. Wang, Y., Nan, X., Zhao, Y., Jiang, L., Wang, H., Zhang, F., Hua, D., Liu, J., Yang, L., Yao, J., & Xiong, B. (2022b). Discrepancies among healthy, subclinical mastitic, and clinical mastitic cows in fecal microbiome and metabolome and serum metabolome. Journal of Dairy Science, 105(9), 7668–7688. 10.3168/jds.2021-21654

56. Wang, Y., Zhao, Y., Nan, X., Wang, Y., Cai, M., Jiang, L., Luo, Q., & Xiong, B. (2022). Rumen-protected glucose supplementation alters fecal microbiota and its metabolic profiles in early lactation dairy cows. Frontiers in Microbiology, 13. 10.3389/fmicb.2022.1034675

57. Xiao, L., Estellé, J., Kiilerich, P., Ramayo-Caldas, Y., Xia, Z., Feng, Q., Liang, S., Pedersen, A., Kjeldsen, N. J., Liu, C., Maguin, E., Doré, J., Pons, N., Le Chatelier, E., Prifti, E., Li, J., Jia, H., Liu, X., Xu, X., … Wang, J. (2016). A reference gene catalogue of the pig gut microbiome. Nature Microbiology, 1. 10.1038/nmicrobiol.2016.161

58. Yang, H., Heirbaut, S., Jing, X., De Neve, N., Vandaele, L., Jeyanathan, J., & Fievez, V. (2022). Susceptibility of dairy cows to subacute ruminal acidosis is reflected in both prepartum and postpartum bacteria as well as odd-and branched-chain fatty acids in feces. Journal of Animal Science and Biotechnology, 13(1). 10.1186/s40104-022-00738-8

59. Zhang, F., Zhao, Y., Wang, Y., Wang, H., Nan, X., Guo, Y., & Xiong, B. (2022). Dietary supplementation with calcium propionate could beneficially alter rectal microbial composition of early lactation dairy cows. Frontiers in Veterinary Science, 9. https://www.frontiersin.org/journals/veterinary-science/articles/10.3389/fvets.2022.940216

60. Zhang, G., Wang, Y., Luo, H., Qiu, W., Zhang, H., Hu, L., Wang, Y., Dong, G., & Guo, G. (2019). The association between inflammaging and age-related changes in the ruminal and fecal microbiota among lactating holstein cows. Frontiers in Microbiology, 10(AUG). 10.3389/fmicb.2019.01803

61. Zhang, J., Jin, W., Jiang, Y., Xie, F., & Mao, S. (2022). Response of Milk Performance, Rumen and Hindgut Microbiome to Dietary Supplementation with Aspergillus oryzae Fermentation Extracts in Dairy Cows. Current Microbiology, 79(4). 10.1007/s00284-022-02790-z

62. Zhao, C., Hu, X., Qiu, M., Bao, L., Wu, K., Meng, X., Zhao, Y., Feng, L., Duan, S., He, Y., Zhang, N., & Fu, Y. (2023). Sialic acid exacerbates gut dysbiosis-associated mastitis through the microbiota-gut-mammary axis by fueling gut microbiota disruption. Microbiome, 11(1). 10.1186/s40168-023-01528-8

63. Zhong, Y., Xue, M. Y., Sun, H. Z., Valencak, T. G., Guan, L. L., & Liu, J. (2020). Rumen and hindgut bacteria are potential indicators for mastitis of mid-lactating holstein dairy cows. Microorganisms, 8(12), 1–13. 10.3390/microorganisms8122042

